# Patient-Derived Orthotopic Xenografts and Cell Lines from Pediatric High-Grade Glioma Recapitulate the Heterogeneity of Histopathology, Molecular Signatures, and Drug Response

**DOI:** 10.1101/2020.12.06.407973

**Authors:** Chen He, Ke Xu, Xiaoyan Zhu, Paige S. Dunphy, Brian Gudenas, Wenwei Lin, Nathaniel Twarog, Laura D. Hover, Chang-Hyuk Kwon, Lawryn H. Kasper, Junyuan Zhang, Xiaoyu Li, James Dalton, Barbara Jonchere, Kimberly S. Mercer, Duane G. Currier, William Caufield, Yingzhe Wang, Alberto Broniscer, Cynthia Wetmore, Santhosh A. Upadhyaya, Ibrahim Qaddoumi, Paul Klimo, Frederick Boop, Amar Gajjar, Jinghui Zhang, Brent A. Orr, Giles W. Robinson, Michelle Monje, Burgess B. Freeman, Martine F. Roussel, Paul A. Northcott, Taosheng Chen, Zoran Rankovic, Gang Wu, Jason Chiang, Christopher L. Tinkle, Anang A. Shelat, Suzanne J. Baker

**Affiliations:** Departments of Developmental Neurobiology, St. Jude Children’s Research Hospital, Memphis TN. 38105; Departments of Center for Applied Bioinformatics, St. Jude Children’s Research Hospital, Memphis TN. 38105; Departments of Oncology, St. Jude Children’s Research Hospital, Memphis TN. 38105; Departments ofChemical Biology and Therapeutics, St. Jude Children’s Research Hospital, Memphis TN. 38105; Departments of Pathology, St. Jude Children’s Research Hospital, Memphis TN. 38105; Departments of Tumor Cell Biology, St. Jude Children’s Research Hospital, Memphis TN. 38105; Departments of Preclinical Pharmacokinetics Shared Resource, St. Jude Children’s Research Hospital, Memphis TN. 38105; Division of Hematology-Oncology, Children’s Hospital of Pittsburgh, Pittsburgh, PA; Exelixis, Alameda, CA; Departments of Surgery, St. Jude Children’s Research Hospital, Memphis TN. 38105; Departments of Computational Biology, St. Jude Children’s Research Hospital, Memphis TN. 38105; Department of Neurology, Stanford University, Stanford, CA; Departments of Radiation Oncology, St. Jude Children’s Research Hospital, Memphis TN. 38105

**Author notes:** Shared first authors contributed equally (Chen He, Ke Xu, Xiaoyan Zhu, Paige S. Dunphy). Shared corresponding authors.

## Abstract

Pediatric high-grade glioma (pHGG) is a major contributor to cancer-related death in children. *In vitro* and *in vivo* disease models reflecting the intimate connection between developmental context and pathogenesis of pHGG are essential to advance understanding and identify therapeutic vulnerabilities. We established 21 patient-derived pHGG orthotopic xenograft (PDOX) models and eight matched cell lines from diverse groups of pHGG. These models recapitulated histopathology, DNA methylation signatures, mutations and gene expression patterns of the patient tumors from which they were derived, and included rare subgroups not well-represented by existing models. We deployed 16 new and existing cell lines for high-throughput screening (HTS). *In vitro* HTS results predicted variable *in vivo* response to inhibitors of PI3K/mTOR and MEK signaling pathways. These unique new models and an online interactive data portal to enable exploration of associated detailed molecular characterization and HTS chemical sensitivity data provide a rich resource for pediatric brain tumor research.

## Introduction

Brain tumors are the predominant cause of cancer-related morbidity and mortality in children^1^. Pediatric diffuse high-grade gliomas (pHGG) comprise approximately 20% of all childhood brain tumors. This heterogeneous group of tumors carries a devastating prognosis, with 70 to 90 percent of patients dying within two years of their diagnosis^1^. Genome-wide analyses have transformed our understanding of pHGGs to illuminate distinct molecular features compared to adult HGG, including a close association between tumor location, patient age, and recurrent mutations that indicates an intimate connection between pHGG pathogenesis and developmental context^2–4^. For example, histone H3 K27M mutations, which are rare in other tumor types, occur in approximately 80% of diffuse intrinsic pontine gliomas (DIPG) and other diffuse HGGs in midline structures such as the thalamus^5–8^. This striking association has redefined the diagnosis of these tumors, with the 2016 World Health Organization Classification of Tumors of the Central Nervous System (CNS) now incorporating molecular-based criteria to define diffuse midline glioma - H3K27M mutant (DMG-K27M) as a distinct diagnostic entity^9^. H3K27M mutations most frequently occur in two of the fifteen genes that encode histone H3, with *H3F3A* encoding H3.3 K27M and *HIST1H3B* encoding H3.1 K27M in approximately 75% and 25% of H3 mutant DIPG, respectively^2^. Activating mutations in the gene encoding the BMP receptor ACVR1 are found almost exclusively in DIPG and preferentially co-occur with H3.1 K27M mutations, generally in younger patients, demonstrating an even more restricted association with developmental context^6,10–13^. In contrast, Histone H3.3 G34R/V mutations are found in approximately 15% of cortical HGG, with patient age ranging from older adolescents through young adulthood^6–8^.

Variable combinations of additional mutations also contribute to intertumoral heterogeneity, including mutations activating the receptor tyrosine kinase-Ras-PI3-kinase pathway, alterations inactivating tumor suppressors *TP53* or *CDKN2A,* mutations in epigenetic regulators such as ATRX, and others^6,7,10–13^. Different subclasses of pHGG are readily detected through comparisons of genome-wide DNA methylation profiles, which may reflect both the developmental origins of the tumors and the consequences of tumorigenic mutations^14^. This epigenetic characterization allows a refined molecular classification of CNS tumors and is increasingly being incorporated into clinical practice.

Despite rapid advances in the characterization of the genomic and epigenetic landscape of pHGG, effective therapeutic approaches are still lacking for almost all pHGG patients^3,15^. *In vitro and in vivo* models that recapitulate the complexity and heterogeneity of pHGG are essential to advance our understanding and identify therapeutic vulnerabilities of these deadly childhood brain tumors.

Here, we report the establishment of a unique collection of 21 patient-derived orthotopic xenograft (PDOX) models and eight new pHGG cell lines recapitulating molecular signatures of the primary tumors from which they were derived and representing a broad spectrum of the heterogeneity found in pHGG. We used a total of 14 pHGG cell lines and two control cell lines for high-throughput screening (HTS) to identify drug sensitivity and validated the *in vitro* heterogeneity of response for two drugs *in vivo.* Detailed molecular characterization of these novel models and the results from HTS chemical sensitivity studies on the large cell line panel are available through an interactive online data portal providing a rich resource to the pediatric brain tumor research community.

## Results

### Patient-Derived Orthotopic Xenografts and Cell Lines of pHGG Represent Diverse Tumor Subtypes

We generated a unique resource of 21 pHGG PDOX by implanting dissociated tumor cells from surgical or autopsy samples into the brains of immunodeficient mice. Patient age at diagnosis ranged from 4-19 years, with a median age of 12 years and median survival of 12 months (Supplementary Table 1). Engraftment efficiency was higher for surgical samples (56%) than for autopsy samples (33%), likely due to decreased tumor cell viability in postmortem material. After initial tumor engraftment, tumors were collected, dissociated, and passaged by intracranial implantation into additional immunodeficient mouse hosts to confirm that the PDOX could be reliably maintained, and for expansion and banking. Passaged tumors were dissected and cryopreserved as viable cells, snap-frozen for subsequent molecular analyses, or processed for histopathology evaluation. The majority of PDOX models were transduced with lentivirus expressing luciferase-2a-YFP to enable *in vivo* imaging^16^. The mouse host survival times for this collection of PDOX models ranged from 2-6 months after intracranial implantation. (Supplementary Table 1). For a subset of the tumors, dissociated cells were adapted for *in vitro* propagation under neural stem cell conditions to facilitate the use of these as matched cell line models for high-throughput drug screening (Fig.1).

**Figure 1:**
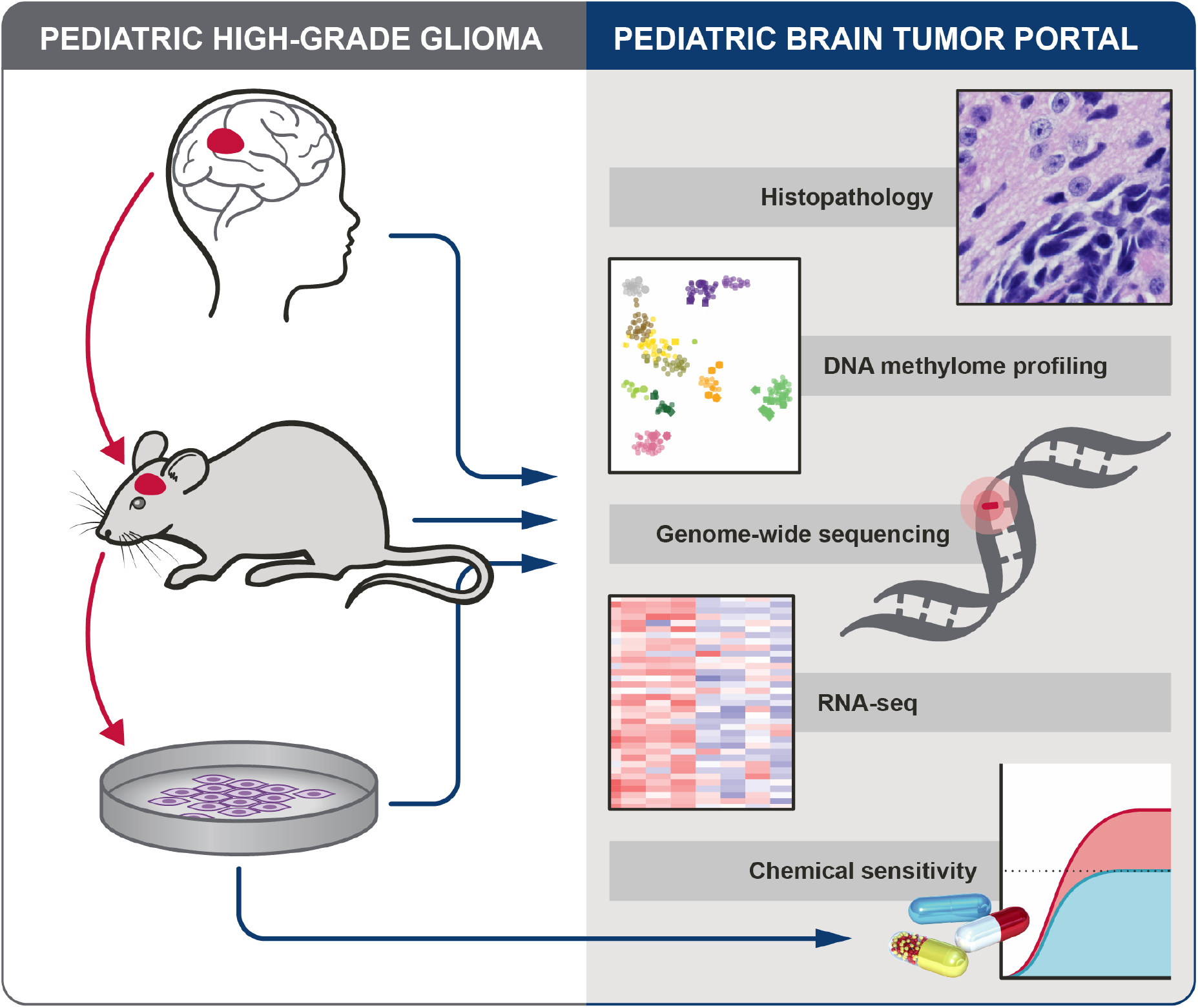
Overview of PDOX and cell line establishment, characterization and preclinical testing, and pHGG data available in Pediatric Brain Tumor Portal.

We performed DNA methylome profiling to classify all 21 PDOX tumors, 19 matched patient tumors from which PDOXs were established, and eight matched cell lines established from PDOXs (Supplementary Table 1b). Using t-distributed stochastic neighbor embedding (t-SNE) analysis of methylation profiles from these samples and a reference set including samples from diverse brain tumor subgroups^14^, all PDOX models and cell lines clearly grouped with glioblastoma or glioma, distinct from embryonal or ependymal tumors. (Fig. 2a). An expanded view of the glioblastoma and glioma clusters shows that 17 of 19 PDOXs and all eight cell lines cluster closely with the tumor from which they were derived (Fig. 2b). Following the established classification scheme^14^, this novel collection of PDOX models comprises six DMG K27M, two pleomorphic xanthoastrocytomas (PXA), 10 glioblastoma, IDH wildtype, including three H3.3 G34 mutant (GBM G34), four subclass midline (GBM Mid), one subclass RTK II (pedRTKII), and two subclass RTKIII (pedRTKIII). Three tumors did not directly match the reference clusters, but were closest to GBM Mid, pedRTKII, or PXA (Fig. 2 and Supplementary Table 1a and b). For two models, SJ-HGGX51 and SJ-HGGX78, the diagnostic patient tumor from which the PDOX was established clustered with reference samples from control inflammatory cells (CONTR_INFLAM), suggesting the presence of large areas of necrosis in the tumor surgical specimens with exuberant inflammatory infiltration, while the PDOX clustered with, or very close to the pedRTKII glioblastoma subgroup. Three PDOX models, SJ-HGGX56, SJ-HGGX58, and SJ-HGGX59 that were established from patient tumors with mismatch repair deficiency (MMRD) clustered tightly with their matched patient tumors, and close to one another, along with midline tumors despite their cortical location in the patients (Fig. 2b). In addition to the eight cell lines established from these PDOX models, we also evaluated methylation profiles from six previously reported DIPG cell lines^17^ used in our preclinical testing experiments. All H3K27M mutant tumors, including patient tumors, PDOXs and cell lines, clustered with the DMG K27M subgroup as expected (Fig. 2a,b).

**Figure 2:**
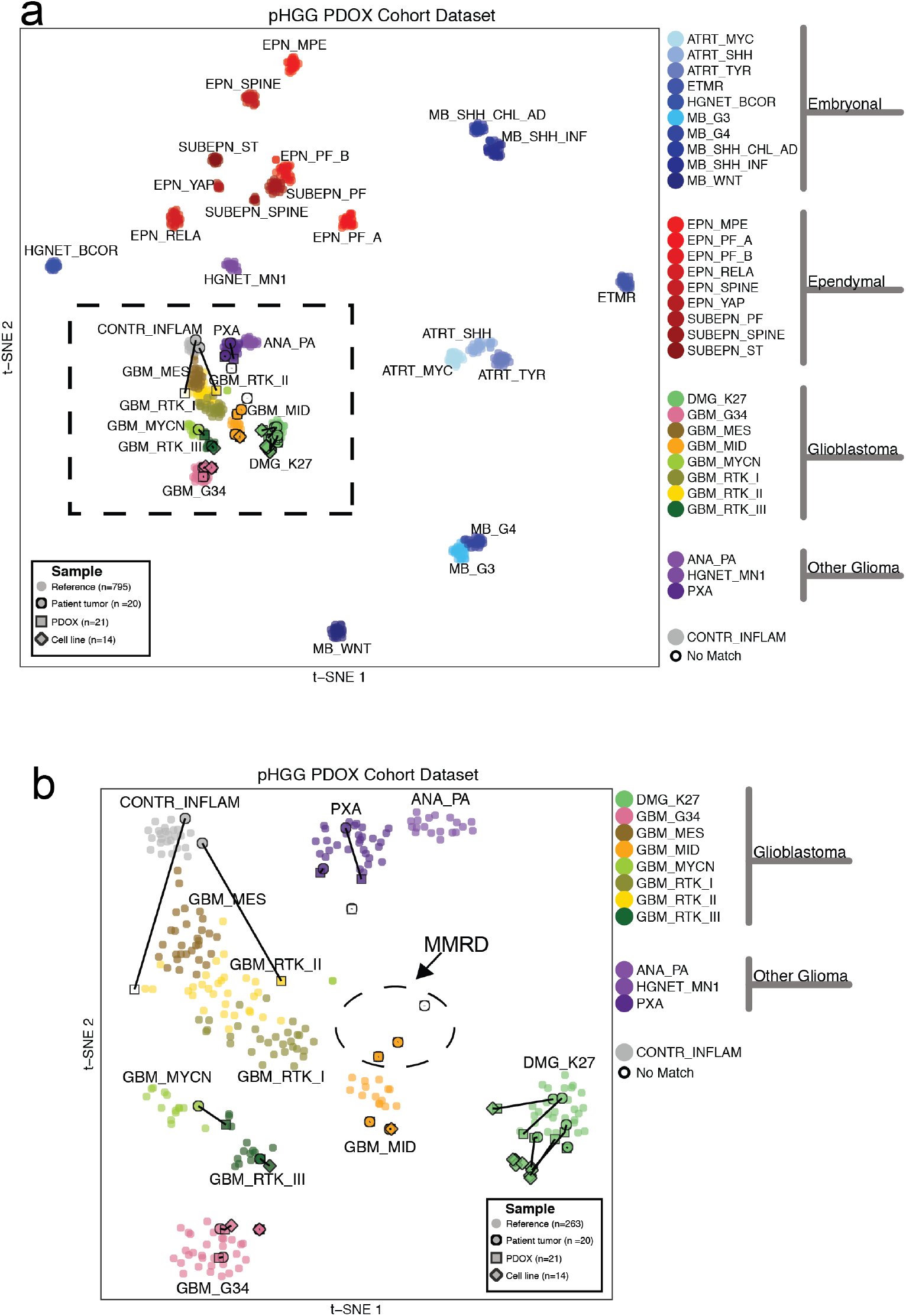
DNA methylation classification of patient tumors is conserved in corresponding PDOXs and cell lines. (a) t-SNE plot showing tumor subgroups based on DNA methylation profiling. Nineteen patient tumors (circles), 21 PDOXs (squares) and 14 cell lines (diamonds) are outlined in black. Lines connect PDOXs and cell lines with the patient tumors from which they were derived. Tumor subgroup classifications are color-coded and circles without outlines are reference samples from Capper et al. Dashed square shows region containing all HGG samples. Classifications: Embryonal Tumors: atypical teratoid rhabdoid tumors (ATRT), embryonal tumor with multilayered rosettes (ETMR), high-grade neuroepithelial tumor with *BCOR* alteration (HGNET_BCOR) and medulloblastoma (MB). Ependymal Tumors: ependymoma (EPN), subependymoma (SUBPEN), myxopapillary ependymoma (MPE), posterior fossa (PF), supratentorial (ST). Glioblastoma: diffuse midline glioma (DMG) and glioblastoma (GBM). Other Glioma: anaplastic pilocytic astrocytoma (ANA_PA), high-grade neuroepithelial tumor with *MN1* alteration (HGNET_MN1), anaplastic pleomorphic xanthoastrocytoma (PXA). Control tissue, inflammatory tumor microenvironment (CONTR_INFLAM). (b) Enlarged view of boxed region in. Dashed oval shows patient tumors with MMRD and derived PDOX models.

Histopathology of PDOX models showed typical characteristics of pediatric HGG including high cellularity, varying extent of astrocytic differentiation, readily apparent mitotic activity, infiltration of the CNS parenchyma, vascular endothelial proliferation and areas of necrosis. PDOX models recapitulated architectural and cytologic features seen in the corresponding patient tumors from which they were derived (Fig. 3).

**Figure 3:**
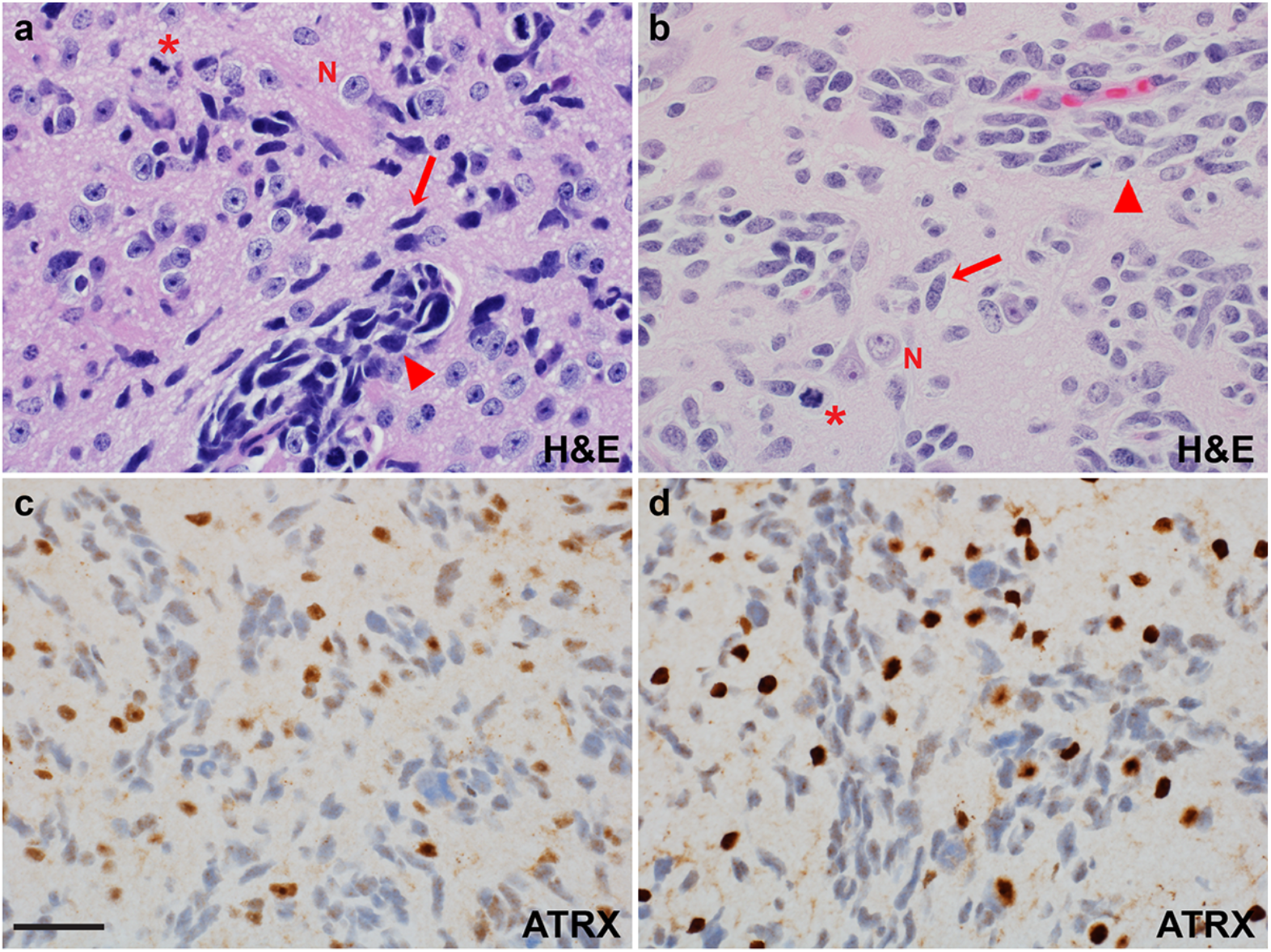
Histopathology of PDOX recapitulates salient features of the patient tumor from which it was derived. H&E staining of HGG PDOX SJ-HGGX6 (a), shown as a representative example, recapitulates histologic features of its corresponding primary human tumor (b), including infiltration of the CNS parenchyma (→), perivascular invasion (►), and apparent mitotic activity (*). Nuclear ATRX immunoreactivity, while retained in the entrapped neurons, is lost in the PDOX tumor cells (c), as in its corresponding primary human tumor (d). N: Entrapped cerebral cortical neurons. Scale bar: 50 μM.

### PDOX and cell lines recapitulate recurrent mutations and gene expression signatures characteristic of pHGG

For a more comprehensive view of the genomic landscape of these models, we performed whole-genome or whole-exome sequencing on all PDOX, matched patient tumors and derived cell lines. Somatic mutations and potentially pathogenic germline mutations were identified for tumors with matched germline samples for 16/21 lines, and potentially pathogenic non-silent mutations were annotated for tumors without available matched germline (Fig. 4, Supplementary Table 2). As expected, there was a dramatically increased mutation burden in PDOXs with MMRD (SJ-HGGX56, 58 and 59) compared to the rest of the cohort (median non-silent SNVs in the exome of 19,336 compared to 26). Recurrent mutations characteristic of pHGG were well-represented in this cohort of tumors, including hotspot mutations in histone H3, as well as mutations in genes involved in chromatin and transcription regulation such as *ATRX, BCOR* and *MYCN,* recurrent alterations in the receptor tyrosine kinase-RAS-PI3-Kinase pathway including missense mutation, amplification and gene fusion of *PDGFRA,* alterations in the *TP53* and RB/cell cycle pathways and activating missense mutation in *ACVR1* (Fig. 4). The great majority of PDOXs and cell lines maintained signature mutations and copy number abnormalities (CNAs) found in the matched patient tumors including large scale gains and losses and focal amplifications in extra-chromosomal DNA, although there were some examples of divergence between patient tumor and derivative models (Fig. 4, Supplementary Fig. 1 and 2). PDOX models also maintained mutations of signature glioma genes over multiple passages, as shown for passages 7 and 10 of SJ-DIPGX7 and passages 3 and 4 of SJ-DIPGX29 (Fig. 4).

**Figure 4.**
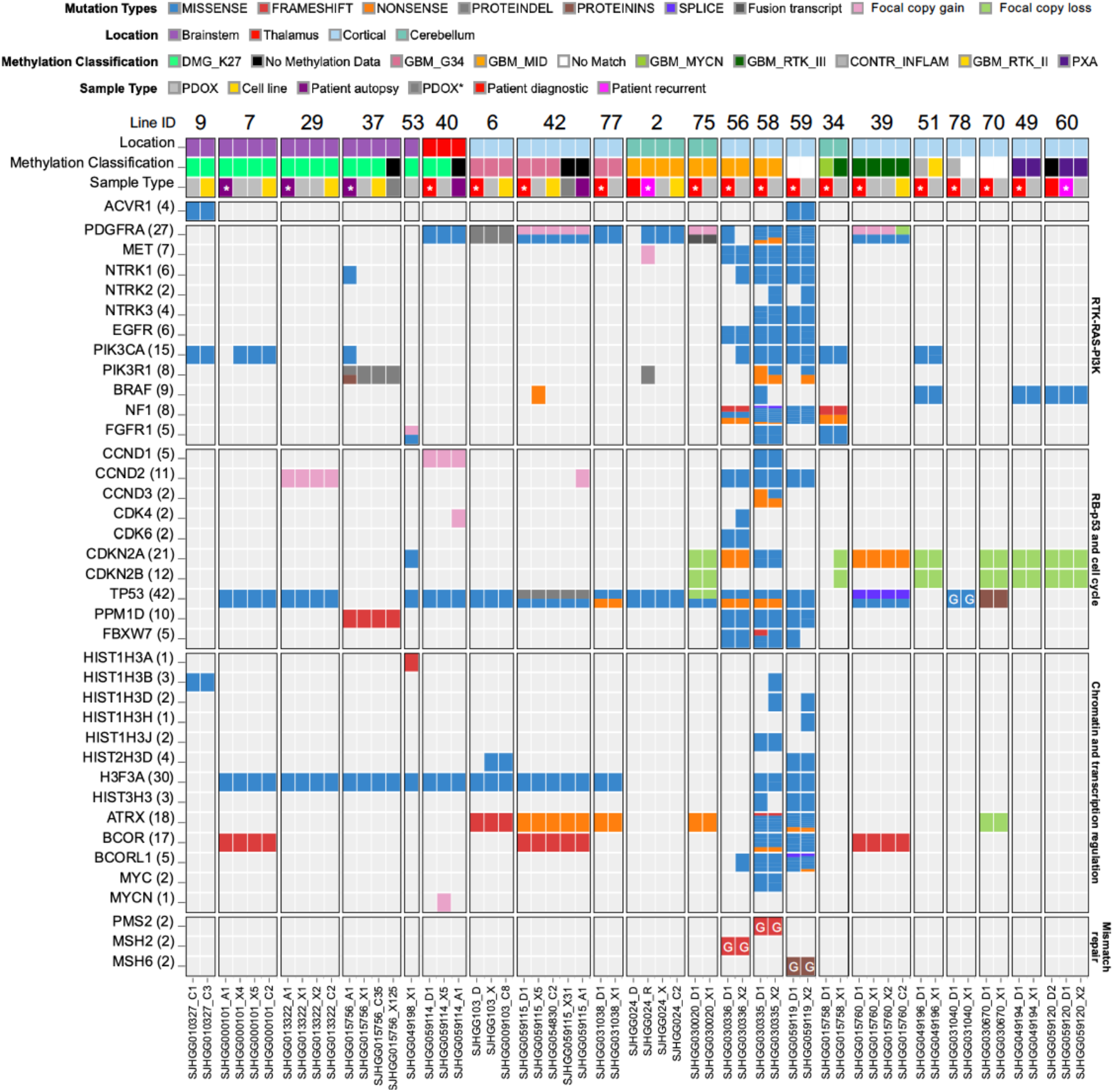
Genomic landscapes of PDOX and cell lines conserve alterations present in the matched patient tumor and represent a variety of pHGG subtypes. Alterations in genes recurrently mutated in pHGG are indicated on the left. Pathways are indicated on the right. Columns show tumor samples. PDOX and cell lines are grouped together with patient tumors from which they were derived. Numbers are the PDOX identifier IDs across the top, sequence file IDs across the bottom. Rows show the location of patient tumors, DNA methylome classification, and tumor sample type. In some cases, patient tumor samples from recurrence or autopsy are included along with the diagnostic sample. Asterisk in tumor sample type indicates the patient tumor from which the PDOX was derived. PDOX* (dark gray box) indicates xenograft generated by implanting the associated cell line. Mutations in signature genes shown as rows are indicated by Mutation Type color code. G in block indicates germline mutation.

We previously showed that pHGG, including DIPG, showed heterogeneous expression signatures recapitulating the glioma subgroups Proneural, Proliferative, and Mesenchymal^17–20^. Analysis of these expression signatures in the entire cohort of patient tumors, PDOXs, and cell lines showed PDOX models represented in all three expression subgroups. However, proliferative signatures were much stronger in general in PDOX and cell line models compared with patient tumors (Fig. 5). Consistent with this observation, analysis of genes that were differentially expressed between PDOX and patient samples across the entire matched cohort (|logFC| >1, adj.p<0.05), showed upregulation of genes associated with cell cycle progression (adj p<2.2E-16), and downregulation of genes associated with inflammatory response (adj p<2.2E-16) in PDOX (Supplementary Table 3). We removed the shared differences between PDOX and patient tumors to better compare the similarity in expression signatures between matched PDOX and patient tumors and found that expression signatures of PDOX correlated strongly with their matched patient tumor (Supplementary Fig. 3). Representative PDOX models retained fidelity of transcriptome signatures over multiple passages (Pearson correlation 0.98, p<2.2e-16), and cell lines, which represent extensive passaging in neural stem cell growth media also showed strong fidelity with the matched PDOX models from which they were derived (Pearson correlation from 0.87-0.95, p<2.2e-16) (Supplementary Fig. 4).

**Figure 5:**
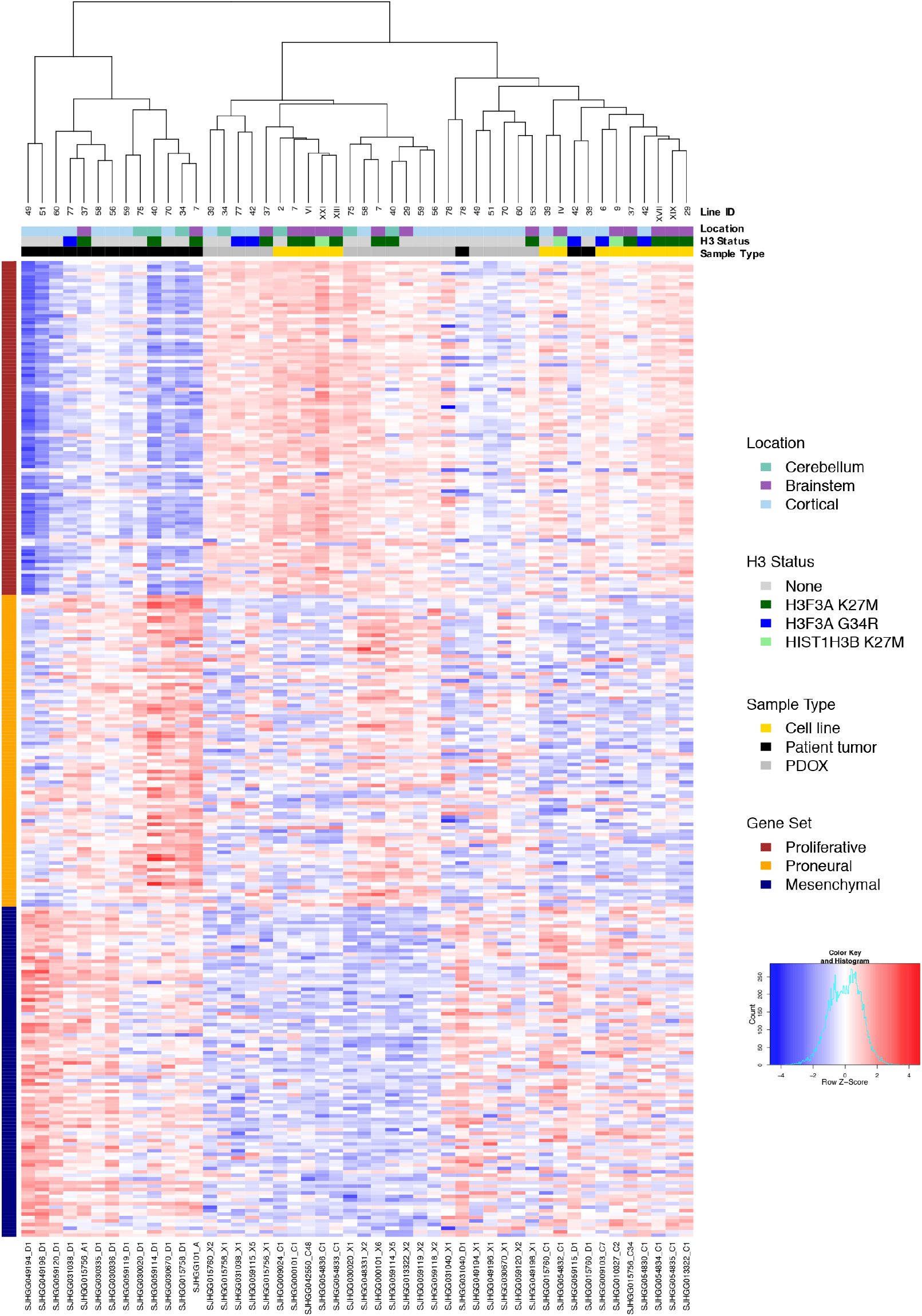
Gene expression signatures of PDOX models recapitulate glioma expression subgroups. Unsupervised hierarchical clustering of RNA-seq quantification (logCPM) of genes from three expression signatures recapitulating glioma subgroups Proliferative, Proneural, and Mesenchymal across the patient tumors, PDOXs, and cell lines.

### Online interactive portal of PDOX and cell line characterization

To maximize the utility of these well-characterized models for the cancer research community, we developed an online Pediatric Brain Tumor Portal (https://pbtp.stjude.cloud) that supports interactive exploration of all molecular data. Within this rich open-access portal, detailed characterization sheets can be viewed or downloaded for each individual model, summarizing clinical, molecular and histopathological features (Supplementary Fig. 5). In addition, this portal will house information about other pediatric brain tumor PDOX models derived at St. Jude Children’s Research Hospital^21^ and will be continually updated with new models in the future.

### High-throughput screen for drug response of different pHGG subtypes

We selected a panel of 16 cell lines to assess drug sensitivity of different subgroups of pHGG: eight glioma stem-cell (GSC) lines were established from PDOXs reported here, six were previously reported patient-derived DIPG cell lines^22,23^ (Supplementary Table 4). Two different sources of proliferating normal human astrocytes were used as controls: astrocytes generated from iPS cells (iAstro), and astrocytes isolated from human embryonic brainstem (HABS) (Supplementary Table 4). Together, the cohort comprised two cortical HGG with H3.3 G34R mutation, two HGG with wildtype histone H3, and 10 DIPG; seven with H3.3 K27M and three with H3.1 K27M mutation. We optimized all cells to grow in 384-well format plates for HTS and conducted a series of validation experiments to ensure reproducibility, including cell proliferation curves, culture conditions, and the automation parameters of HTS. We showed that DIPG cells plated as adherent cultures on basement membrane matrix compared with the same cells grown as tumorspheres responded similarly to a panel of 53 different drugs representing a range of mechanisms of action (MOA) tested in a dose-response (DR) format (Supplementary Fig. 6, Pearson correlation = 0.994; Supplementary Table 5 a and b). Therefore, we conducted HTS experiments with adherent cells to avoid variability within and between cultures introduced by variable tumorsphere sizes.

As an initial screen for sensitivity to drugs that could be quickly deployed in the clinic, we tested 1,134 FDA-approved drugs at a single concentration in nine pHGG cell lines (SJHGGx39c, SJHGGx6c, SJDIPGx37c, SJDIPGx29c, SJDIPGx9c, SJDIPGx7c, SUDIPG-XIII, SUDIPG-IV, SUDIPG-VI) and human embryonic stem cell-derived neural stem cells (hNSC) as a normal cell type reference. Top hits from this screen were supplemented with recently FDA-approved oncology drugs, epigenetic modulators, clinical candidates, and relevant chemical probes to yield a set of 93 drugs that were screened against the full panel of 16 cell lines in DR format using an independently prepared compound plate. For all 16 cell lines, the z-prime values^24^ calculated from 310 384-well assay plates showed an excellent signal to noise ratio (z-prime values: median=0.82, minimum=0.56, maximum=0.92) (Supplementary Fig. 7). An additional 246 compounds, sampling a broader range of drug mechanisms of action, were screened in DR format in 4 exemplar pHGG models representing different histone sub-types: SJHGGx39c (wild-type histone H3), SJHGGx6c (H3.3 G34R), SJDIPGx29c (H3.3 K27M), and SJDIPGx37c (H3.3 K27M). Results from all stages of these HTS studies are presented in Supplementary Table 5 and are available online for exploration in the Pediatric Brain Tumor Portal (https://pbtp.stiude.cloud) where interactive features allow the user to query by drug class, specific compound name, tumor subgroup, or tumor cell lines, and to visualize results in multiple formats including data on the range of responses to selected compounds, the sensitivity to all tested drugs for selected cells lines, and customized overlay dose-response curves. A mouse over feature also allows the user to identify outliers and view compound information, dose response values and the associated dose response curve.

Results from the comprehensive screen of 93 compounds in 16 cells lines are summarized in Fig. 6. To highlight the most effective drug for each cell line in terms of selectivity, we calculated the area under the curve (AUC) from the DR of each drug and subtracted the median AUC for that drug calculated over all cell lines (Fig. 6a). Notably, the distribution of responses for SUDIPG-XXI showed that these cells were more sensitive than other lines to nearly every drug tested. Consequently, we flagged this model as an outlier and removed it from subsequent analyses. Several drugs were identified as the most selective for more than one pHGG model: the MEK inhibitor trametinib for SJDIPGx9c and SUDIPG-XIX, the NF-κB inhibitor EVP4593 for SJDIPGx7c and SUDIPG-XIII, and the HSP90 inhibitor tanespimycin for SJDIPGx37c and SJHGGx39c. The GSK3 inhibitor LY2090314 was the most selective drug for both SUDIPG-IV and HABS control cells, whereas the proteasome inhibitor marizomib was the most selective for SUDIPG-VI and iAstro control cells.

**Figure 6:**
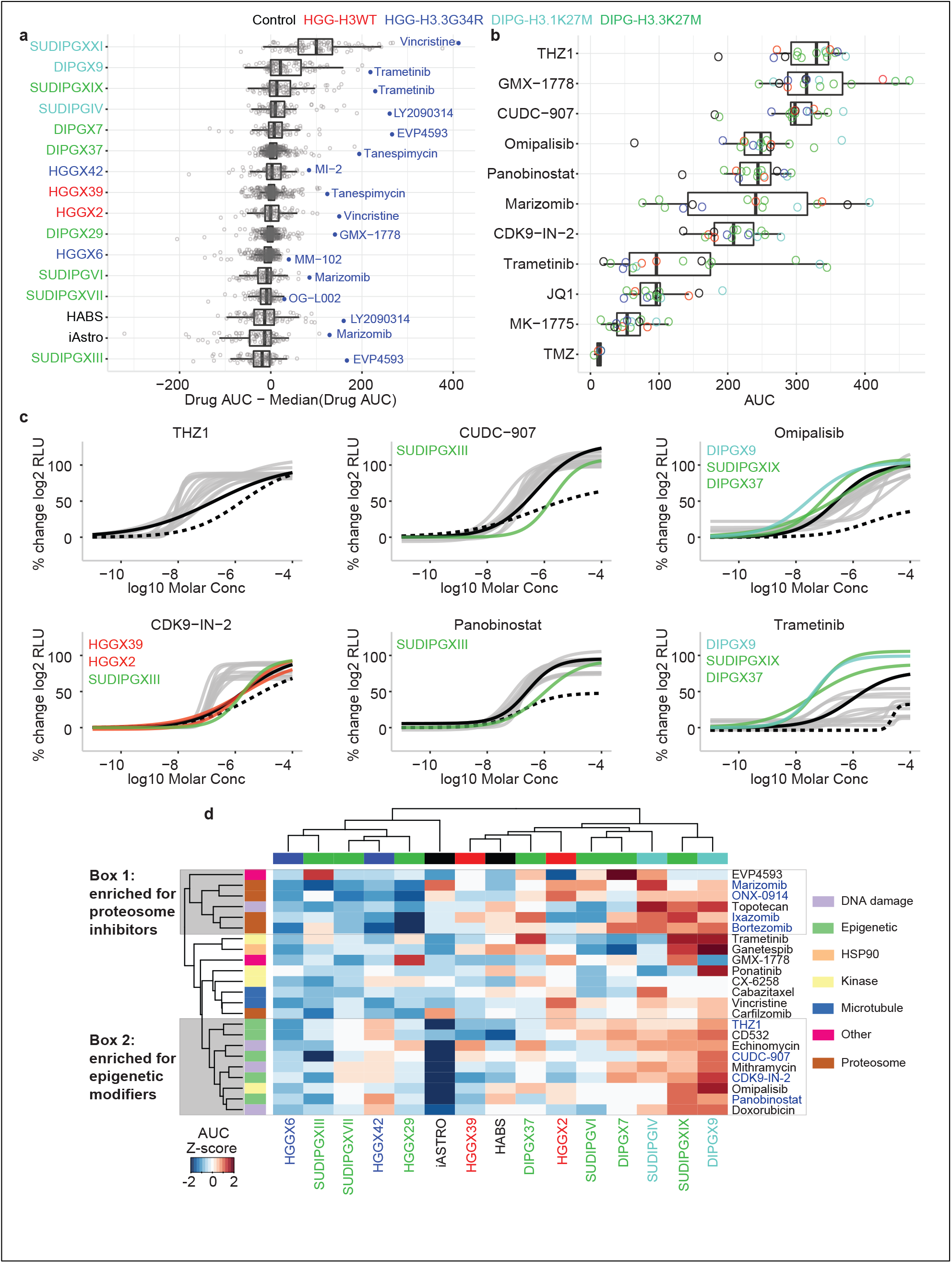
Analysis of screening results from 93 compounds across 14 pHGG models and two normal astrocyte lines. (a) Distribution of normalized drug AUC (the drug AUC in that cell model minus the median AUC for that drug across all models). Each dot represents the result from a single drug tested in the associated model. Normalizing in this way controls for the inherent potency of the drug and emphasizes selectivity across models. The most selective drug for each model is highlighted in blue. Cell models are color coded by histone mutation status. (a)Distribution of AUC for select drugs that have been evaluated in clinical trials or implicated as promising agents in preclinical studies for adult or pediatric glioma. Each dot represents the result for that drug in one model and is color coded by the histone mutation status of the model. (c) Select dose-response curves for the drugs highlighted in (b). Normal reference iAstro is depicted in black dashed lines and HABS in black solid lines. pHGG models are colored gray or by histone mutation status for specific models indicated. (d) Unsupervised hierarchical clustering of drug AUC z-scores for the 25% most active compounds out of the 93 tested. Column and column labels are color coded by histone mutation status. Clusters 1 and 2 (gray boxes) highlight two compound clusters that show distinct activity profiles across the models in this study. The color code for histone mutation status is: H3-wt (red), H3.3 G34R (blue), H3.1 K27M (turquoise), and H3.3 K27M (green). Control cell lines (iAstro and HABS) are black. Color code or mechanism of action is shown on the right and annotated in heatmap row color blocks at left in d.

Trametinib is an FDA-approved drug and we decided to compare the distribution of its AUC values across all 15 models to other drugs in the set that have been evaluated in clinical trials or implicated as promising agents in preclinical studies for adult or pediatric glioma (Fig. 6b-c, Supplementary Fig. 8a). THZ1, which targets CDK7^25^, was the most active drug (highest median AUC) in this subset and was more selective for pHGG models compared to iAstro and HABS. The CDK9 inhibitor CDK9-IN-2^26^ had lower AUC values compared to THZ1, but also showed selectivity for pHGG models relative to control cell lines, although SJHGGx39c, SJHGGx2c, and SUDIPG-XIII were refractory relative to the other tumor models. CDK7 and CDK9 are key regulators of transcription initiation and elongation, respectively, supporting the concept of targeting transcriptional dependencies in tumor cells^27^. In contrast, the pan-BET inhibitor JQ1^28,29^, which also impacts transcription regulation, was much weaker than the CDK7 and CDK9 inhibitors, and was equally or more cytotoxic to control cell lines (Fig. 6b). The anti-metabolite GMX-1778, which disrupts the regeneration of NAD^+^ via NAMPT, showed strong efficacy in all cell lines, in agreement with previous reports in some DIPG models^30^. However, we did not observe enhanced sensitivity in a line with *PPM1D* mutation (SJDIPGx37c) as predicted by previous studies^30^. Consistent with previous reports that evaluated broad-spectrum HDAC *inhibitors^17,22,31,32^,* panobinostat showed strong efficacy *in vitro.* However, CUDC-907, a dual-acting inhibitor of class I PI3K and HDAC^33^, had higher median AUC and was slightly more selective for pHGG models. Both drugs were considerably less active in iAstro but also less effective in SUDIPG-XIII (Fig. 6b,c). The proteasome inhibitor marizomib is being evaluated in a phase I combination study with panobinostat in DIPG (NCT04341311)^17^. This drug had the largest interquartile range of AUC values in the sub-set and induced significant cytotoxicity in iAstro cells (Fig. 6b, Supplementary Fig. 8a). Inhibitors of PI3K/mTOR (omipalisib) and MEK (trametinib) were selective for the same pHGG models over all other cell lines: SJDIPGx9c, SUDIPG-XIX, and SJDIGXc37 (Fig. 6c). However, trametinib was generally much less active in the panel as evidenced by its significantly lower median AUC (Fig. 6b).

The WEE1 inhibitor MK-1775 (adavosertib), which is being evaluated clinically with radiotherapy in pHGG (NCT01922076), showed low efficacy in our panel. Likewise, the alkylating agent temozolomide (TMZ), a standard of care in adult gliomas, was inactive in our tumor models, consistent with the lack of clinical response to TMZ in children with pHGG^34,35^. We also tested two other alkylating agents previously tested in brain tumors, streptozocin and nimustine^36^, and both were inactive in our pHGG models (Supplementary Table 5 c and e).

Finally, to better explore patterns of drug sensitivity across the cell lines in our panel, we performed unsupervised clustering of cell lines (columns) and compounds (rows) with all 93 compounds (Supplementary Fig. 8b), or using the top 25% most active drugs out of the 93 evaluated (Fig. 6d). Notably, tumor models did not cluster according to histone subtype based on their responses to the drugs in this subset. SJHGGx6c and SUDIPG-XIII tended to be more refractory, and SUDIPG-XIX and SJDIPGx9c more sensitive, to this subset. We observed two compound clusters that were enriched for molecules acting by the same mechanism of action. Cluster 1 contained 4 proteosome inhibitors (marizomib, ONX-0914, ixazomib, and bortezomib) and was marked by uniformly weak efficacy against a group of tumors clustered on the left side of the heatmap: SJHGGx6c, SUDIPG-XIII, HGGx42c, and HGGx29c. Cluster 2 contained 4 compounds targeting transcriptional dependencies (THZ1, CDK9-IN-2, panobinostat and CUDC-907) and two dual activity compounds including PI3K inhibition (CUDC-907 and omipalisib), and was characterized by a significant lack of efficacy against the iAstro control cell line, therefore showing some selectivity for HGG.

### Inhibition of PI3K/mTOR and MEK signaling show selective effects alone and in combination in pHGG *in vitro* and *in vivo*

We sought to determine if sensitivity differences between tumor cell lines detected by HTS translated into *in vivo* effects in orthotopic brain tumors. For these studies, we chose PI3K/mTOR and MEK pathway inhibitors because responses in different pHGG lines varied and drug engagement of the target can be reliably detected in tumor tissue. HTS results showed that the PI3K/mTOR inhibitor omipalisib was active in most pHGG models but varied in potency. In contrast, the MEK inhibitor trametinib showed weak activity in many tumor cells (Figure 6 and Supplementary Table 5f). While these pathways are compelling targets for *in vivo* treatment given known genetic aberrations in both pathways in pHGG^2,6^, both compounds are known substrates of ATP-dependent efflux pumps P-gp and BCRP, limiting their brain exposure^37,38^. Therefore, we selected paxalisib (GDC0084) and mirdametinib (PD0325901), which target PI3K/mTOR and MEK, respectively, and have superior blood-brain barrier penetration and exposure^37,39,40^, to validate effects of pathway inhibition for *in vitro/in vivo* comparisons.

Numerous studies have shown extensive cross-talk between these pathways and promise for enhanced efficacy of combined inhibition approaches^41,42^. We conducted *in vitro* quantitative synergy assays in seven cell lines and analyzed results using the Bivariate Response to Additive Interacting Doses (BRAID) response surface model^43^. BRAID κ denotes the type of interaction (κ < 0 is antagonistic; κ = 0 is additive, and κ > 0 is synergistic), whereas BRAID IAE50 computes the degree to which a combination achieves a minimal efficacy within a defined concentration range (in this study, 50% reduction in cell viability for concentrations ≤ 1 μM). Higher IAE50 means that the combination is more efficacious. The combination of paxalisib and mirdametinib exerted synergistic growth inhibition (κ > 0) in three pHGG cell lines (SJ-DIPGX29c, SJ-DIPGX37c and SJ-HGGX6c) and iAstro controls, but not in SJ-DIPGX7c, SJ-HGGX42c, SJ-HGGX6c or HA-bs controls (Figure 7). The drug combination was clearly most efficacious in SJ-DIPGX37c (IAE_50_ = 3.3), followed by SJ-DIPGX29c, SJ-DIPGX7c, SJ-HGGX42c, and then SJ-HGGX6c and the two controls, HA-bs and iAstro. When considering the efficacy of the combination (IAE_50_), synergy made a major contribution in the case of SJDIPGx37c and SJDIPGX29c, as indicated by the curvature of the 50% (black) and 90% (white) cell viability isoboles. In contrast, the IAE_50_ value for SJDIPGx7c was driven solely by paxalisib. Here, the 50% and 90% isoboles run parallel to the y-axis because mirdametinib exerts little cytotoxicity on its own and fails to potentiate the activity of paxalisib. While the synergy is highest in iAstro (κ = 8.8), as evidenced by the clear shift in the 50% isobole with increasing mirdametinib concentrations, both drugs are weakly cytotoxic on their own and their interaction is insufficient to induce high combined efficacy (IAE_50_ = 1.0).

**Figure 7:**
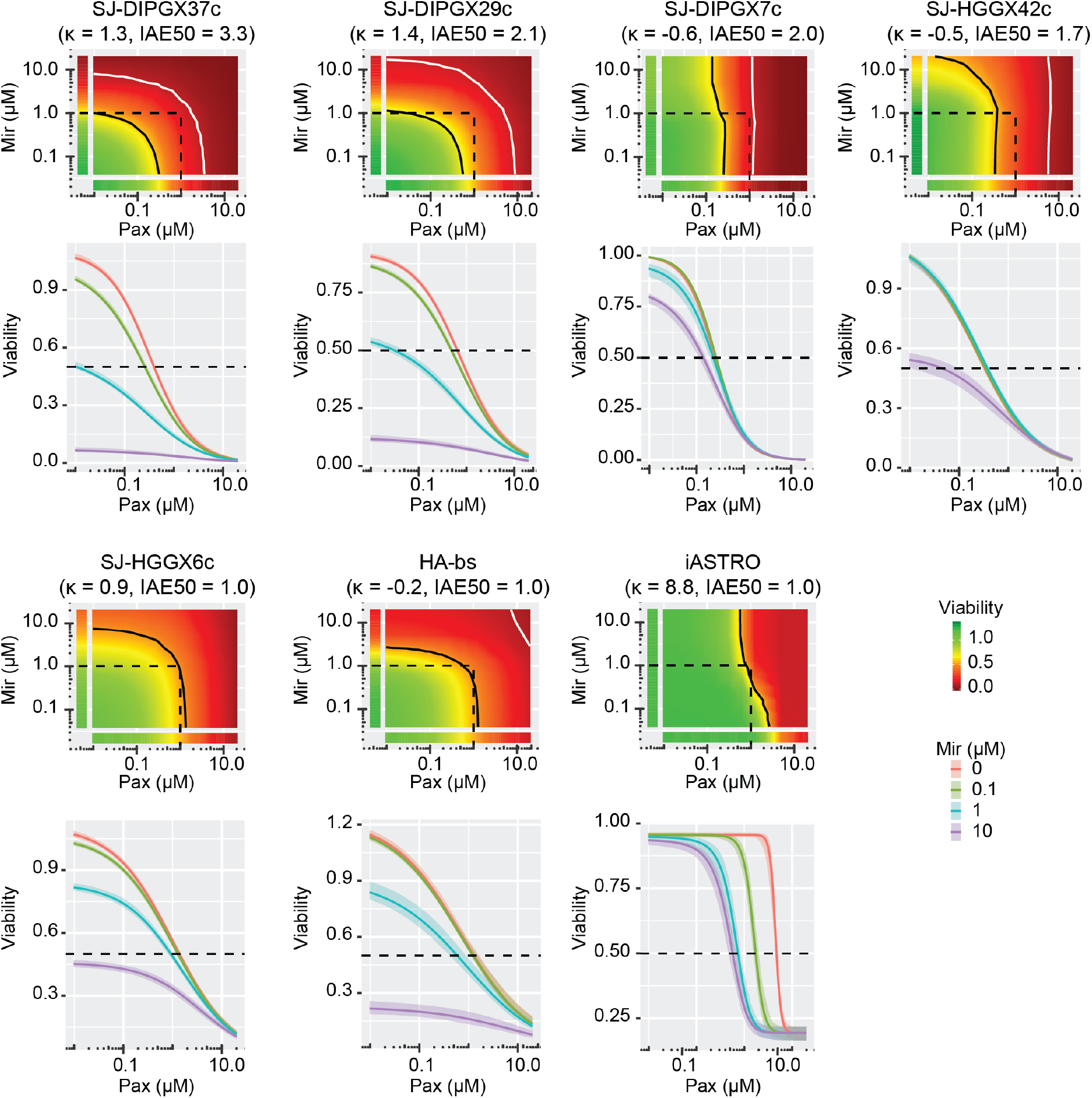
Paxalisib and mirdametinib drive synergistic growth inhibition in a subset of pHGG cell lines. The BRAID model presents synergistic effects of paxalisib (pax) and mirdametinib (mir) following a 7-day treatment in 7-cell lines: H3.3 K27M DIPGs; SJ-DIPGX37c, SJ-DIPGX29c, SJ-DIPGX7c, H3.3 G34R pHGGs; SJ-HGGX42c, SJ-HGGX6c, and astrocyte controls HABS and iAstro. The parameter κ measures the type of interaction: κ<0 implies antagonism, κ = 0 implies additivity, and κ>0 implies synergy). The index of achievable efficacy (IAE) quantifies the degree to which the drug combination achieves a minimal level of efficacy within a defined concentration range. Higher IAE means the combination was more efficacious. In this experiment, it was defined as a 50% reduction of cell viability (black line) at concentrations ≆ 1 μM (dotted lines). The 90% reduction of viability isobole (white line) is included for reference.

We selected two H3.3 K27M mutant DIPG PDOX models, SJ-DIPGX37 and SJ-DIPGX7, to test whether *in vitro* HTS results predicted *in vivo* response. Both harbor mutations targeting PI3K and TP53-related pathways (Fig. 4) and their corresponding cell lines (SJ-DIPX7c and SJ-DIPGX37c) demonstrated different *in vitro* responses. Both cell lines showed similar sensitivity to paxalisib (EC_50_ = 0.32 μM for SJ-DIPGX37c vs. 0.22 μM for SJ-DIPGX7c), whereas mirdametinib induced a stronger response in DIPGX37c (EC_50_ = 1 μM), which has *PIK3R1* and *PPM1D* mutations, and did little in SJ-DIPGX7c, which has *PIK3CA* and *TP53* mutations (Figs. 4 and 7), similar to the responses observed with omipalisib and trametinib in the HTS data (Supplementary Table 5f).

We evaluated PI3K and MEK pathway inhibition in intracranial PDOX models matched to these cell lines by evaluating levels of p-AKT S473 and p-ERK T202/Y204, respectively (Fig. 8 and Supplementary Fig. 9). Interestingly, while the levels of PI3K pathway activity, as assessed by p-AKT, were similar between both PDOXs, SJ-DIPGX7 showed markedly lower levels of p-ERK, indicating lower MEK pathway activation (Fig. 8a). Consistent with a reduced reliance on MEK signaling, mirdametinib did not significantly alter cell survival or proliferation in SJ-DIPGX7, as assessed by active caspase 3 and phospho-histone H3, respectively. Paxalisib treatment alone induced a trend of Increased cell death, and when combined with mirdametinib significantly increased cell death (Figure 8a). SJ-DIPGX37 showed a significant decrease in tumor cell proliferation *in vivo* with paxalisib treatment, and a greater magnitude effect with mirdametinib, while neither induced significant tumor cell death. Strikingly, the combination of paxalisib and mirdametinib significantly enhanced tumor cell death *in vivo* beyond levels induced by either agent alone (Fig. 8b). Consistent with *in vitro* synergy studies, the combination had a much more significant impact on cell death in SJ-DIPGX37.

**Figure 8:**
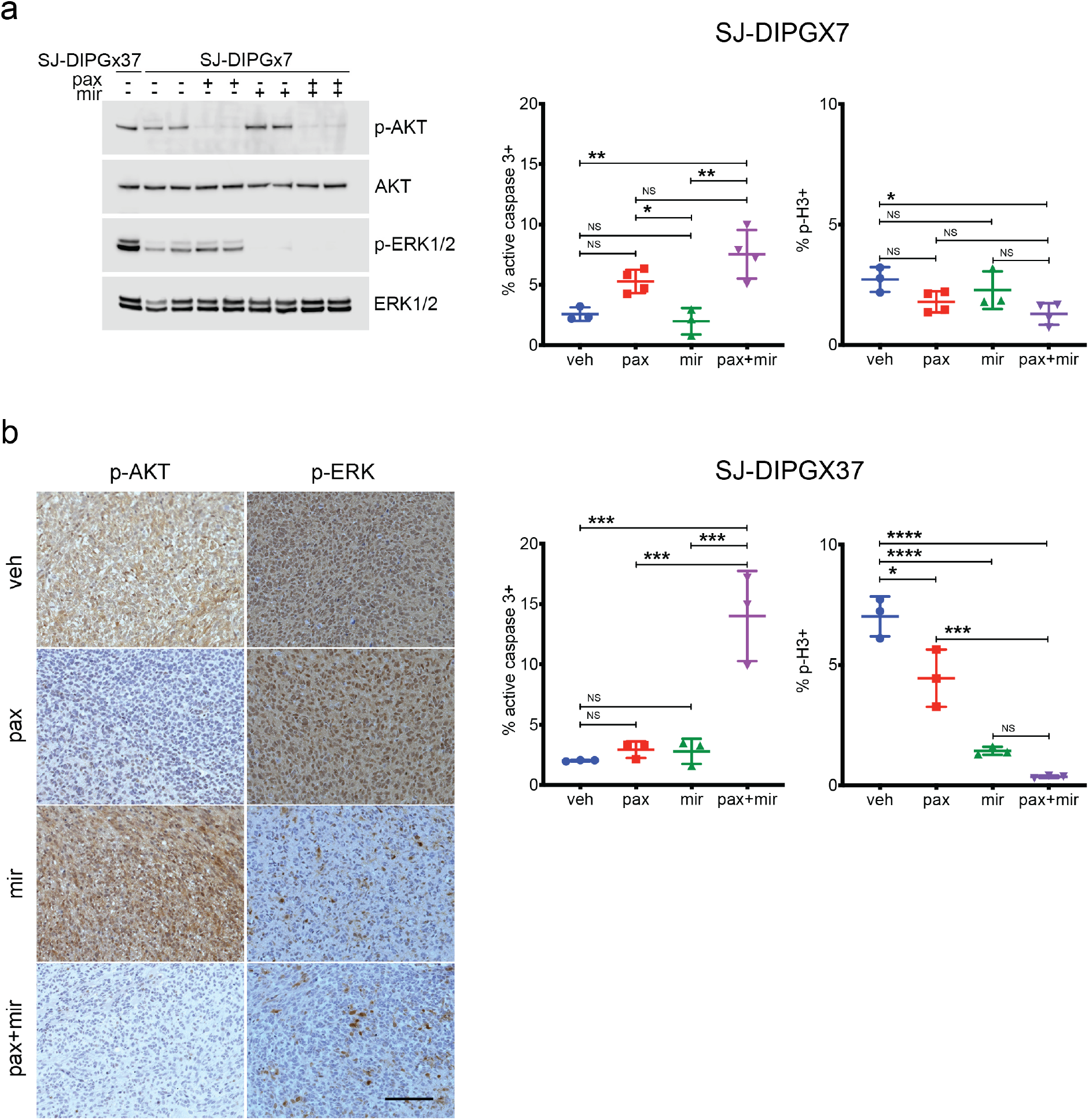
Paxalisib and mirdametinib show selective effects on cell survival and proliferation *in vivo*. (a). Left: Western blot of lysates from intracranial PDOX SJ-DIPGX37 (untreated, lane 1), and SJ-DIPGX7 (lanes 2-9) treated with vehicle (veh), paxalisib (pax), mirdametinib (mir) or the combination of paxalisib and mirdametinib (pax+mir) as indicated; antibodies are indicated at right. Quantification of IHC in sections from SJ-DIPGX7 tumors treated with agents shown along × axis for active caspase 3 (middle) and phospho-histone H3 (right), n= 3 tumors for veh, mir and n=4 tumors for pax and pax+mir. (b). Left: IHC for pAKT Ser473 and pERK in SJ-DIPGX37 tumors in representative tumors treated with veh, pax, mir, or pax+mir as indicated. Quantification of IHC staining in sections from SJ-DIPGX37 tumors treated with agents shown along × axis for active caspase 3 (middle) and phospho-histone H3 (right), n=3 tumors for each treatment. NS, not significant; * p<0.05; **p<0.01; ***p<0.001, ****p<0.0001 using ANOVA with post-hoc Tukey test. Scale bar in B left panel = 100 μM.

We further extended these promising results in SJ-DIPGX37 to evaluate effects on the survival of tumor-bearing mice. We reduced doses of paxalisib to 8mg/kg and mirdametinib to 14mg/kg in monotherapy controls and in the combination for long-term treatment due to significant weight loss. These doses still effectively inhibited both pathways in the brain, with the combination showing slightly enhanced suppression of p-ERK1/2 relative to mirdametinib alone (Supplementary Figure 9c). We randomly distributed 24 mice bearing SJ-DIPGX37 PDOX into four arms; 1) vehicle, 2) paxalisib, 3) mirdametinib, or 4) combination and found that the combination of paxalisib and mirdametinib, but not either drug alone, significantly extended survival of mice with intracranial SJ-DIPGX37 tumors (Fig. 9, p=0.0056, Mantel-Cox log-rank test). We examined the plasma and brain pharmacokinetics of mirdametinib and paxalisib in normal mice to assess the possibility that the enhanced combinatorial effects were due to drug-drug interaction. While both agents had slightly increased plasma AUCs in combination (Supplementary Fig. 10), the differences were within variability and less than twofold magnitude, the conventionally accepted threshold to identify drug-drug interactions^44,45^. Thus, the enhanced efficacy with combination treatment was likely due to combined pathway inhibition.

**Figure 9:**
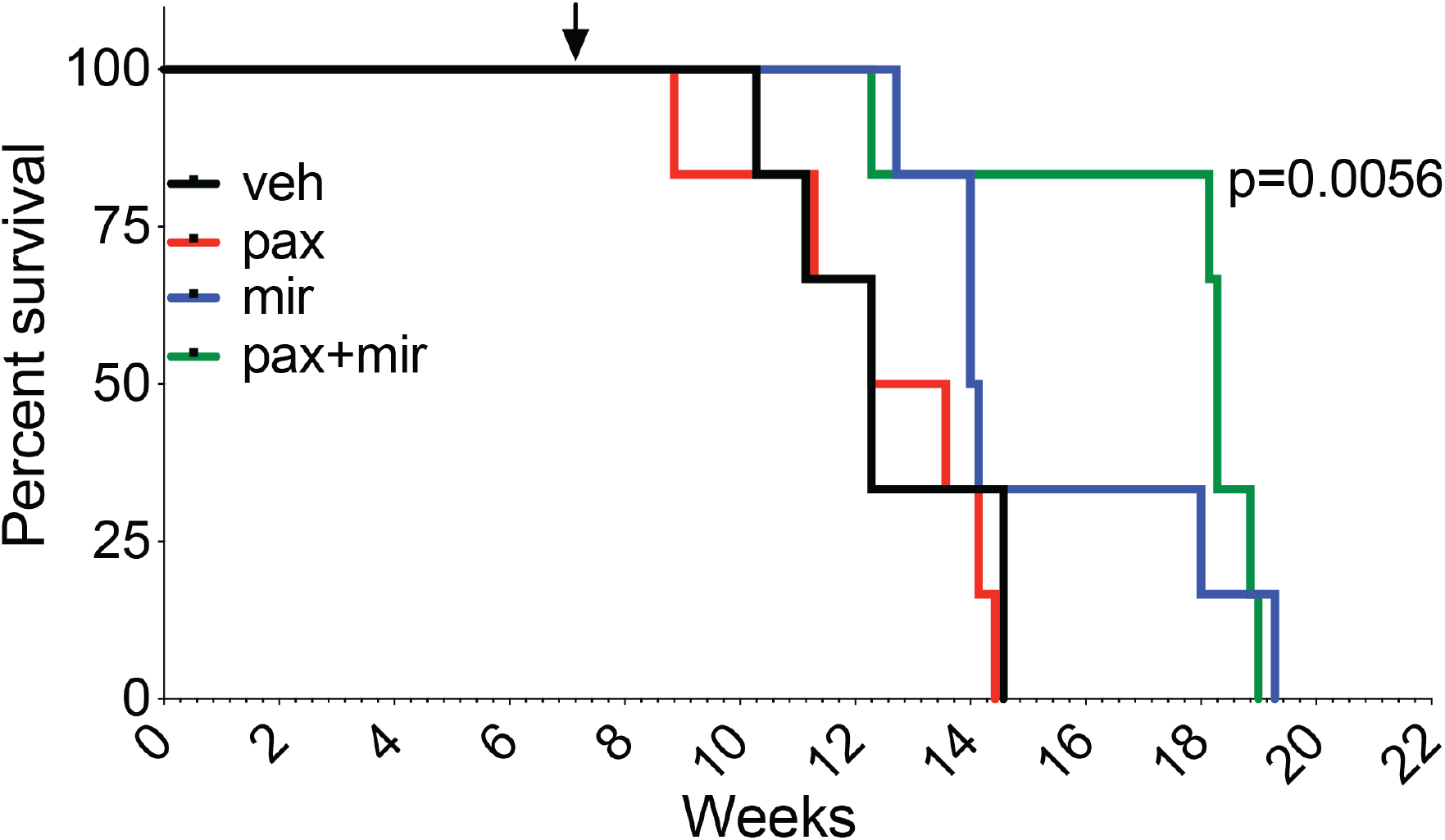
Combined treatment with paxalisib and mirdametinib significantly extends survival of SJDIPGx37 bearing mice. Mice were randomized 50 days after implantation into 4 treatment arms (6 mice per arm) and treated with vehicle, paxalisib (8 mg/kg), mirdametinib (14 mg/kg), or paxalisib + mirdametinib daily, 5-days ON and 2/3-days OFF. paxalisib + mirdametinib vs vehicle, p=0.0056, mirdametinib vs vehicle, p=0.16, paxalisib vs vehicle, p=0.48 (Mantel Cox log-rank test). Arrow shows the time point for randomization and initiation of treatment. Kaplan-Meier survival analysis.

## Discussion

Relevant *in vitro* and *in vivo* disease models that are founded in the correct developmental origins, recapitulate genetic and epigenetic signatures and represent the significant heterogeneity of pHGG are essential to further our understanding of mechanisms driving tumorigenesis and to identify therapeutic vulnerabilities. Although a growing number of DIPG cell lines have been established and characterized^17,22,46–48^, relatively few of these efficiently engraft in the brain as xenografts^49,50^, and there are a much smaller number of cell lines from pediatric gliomas arising outside the brainstem. Studying PDOXs in addition to cell lines allows researchers to address important dimensions of tumor biology, including angiogenesis, tumor invasion, and interactions with the tumor microenvironment that may strongly influence tumor growth and selective pressures, including the contribution of nervous system activity in driving glioblastoma and DIPG growth^51^. A recent study to establish a biobank of pediatric brain tumors reported the establishment of 8 new pHGG PDOX models^52^. The 21 new pHGG PDOX models and eight new cell lines reported here are a significant advance in available pHGG models and include several rare tumor subtypes for which models are currently extremely limited, including three H3.3 G34R glioblastomas, three pHGG models with MMRD, two glioblastomas in the pedRTKIII subgroup, and two PXAs. This new collection of models recapitulates the histopathological and molecular hallmarks of pHGG and preserves mutation and DNA methylome signatures of the primary tumors from which they were derived, including single-nucleotide variants, large scale copy number changes, and the presence of double minute chromosomes.

pHGG is known to display significant intra-tumoral heterogeneity due to clonal variation within the tumor^53–55^. Indeed, there were a few differences between patient tumor, PDOX and cell line, such as *NTRK1* mutation detected in the autopsy sample of SJ-DIPGX37, but not the associated PDOX or cell line, or *MYCN* amplification present in SJ-DMGX40 PDOX, but not detected in the matched patient tumor (Fig. 4, Supplementary Fig. 2). Such differences could be due to regional heterogeneity between the patient sample analyzed compared with cells implanted, expansion of a minor subclone during PDOX establishment, or acquisition of a new mutation during ongoing tumor evolution in the mouse host. Across multiple PDOX models, we found consistent upregulation in genes associated with proliferation and decreased expression of genes associated with inflammatory response compared to matched patient tumors, as previously described^52^. After removing these shared PDOX-dependent signatures, however, there was a strong correlation in expression signatures between PDOX and matched patient tumors.

Despite decades of clinical trials, no effective chemotherapy approaches have been identified for pHGG. However, the revolution in knowledge gained through genomic analyses of pHGG revealed some clear therapeutic targets, and selective inhibitors have induced striking responses in pHGG with *BRAF* V600E mutation or *NTRK* fusion genes^56^. The high frequency of H3 K27M mutations in DIPG and other midline gliomas sparked an intense search for selective vulnerabilities conferred by this mutation, especially in agents connected to epigenetic regulation. Some of the top hits from our HTS were consistent with previous findings in DIPG cell line screening, including inhibitors of HDACs, CDK7 and CDK9, and proteins intersecting with epigenetic regulatory mechanisms and converging on transcription regulation^17,22,27,31,57–59^. Challenges in achieving effective concentrations of the HDAC inhibitor panobinostat in the brain has led to investigations identifying combination approaches to enhance efficacy^17,27,31,32,60,61^. Overall, two sources of normal astrocyte control cells were among the least sensitive cell lines to the collection of compounds tested in DR (Fig. 6a), and many of the top hits showed greater sensitivity in tumor cell lines than in iAstro control cells (Fig 6b and c). However, responses varied between the two sources of astrocytes (Fig. 6b,c and Supplementary Fig. 8). These results reveal potential differences in drug sensitivity in normal cells at different developmental stages as well as variation among tumors, and further highlight the difficulty in determining simple predictors of drug responsiveness in pHGG which are heterogenous in developmental expression signatures as well as genetic mutations.

Understanding how disease heterogeneity contributes to therapeutic response is the foundation of precision medicine. The panel of 14 pHGG cell lines and two astrocyte controls revealed substantial variation in efficacy and potency across the collection of compounds representing multiple MOA. We further investigated inhibitors of PI3K/mTOR and MEK pathways in two PDOX models with differing responses as test cases to determine how well results of *in vitro* screening were predictive of *in vivo* response. Myriad mutations in pHGG and other cancers converge on these two signaling pathways, and there are multiple examples of pathway cross-talk that compromise the efficacy of single-agent approaches that inhibit only one of these two regulatory cascades^41,42,62–64^. Among available inhibitors of PI3K/mTOR and MEK, we selected paxalisib and mirdametinib, respectively, for *in vivo* studies because of their ability to traverse the blood-brain barrier. *In vivo* testing with these drugs induced dramatic inhibition of pathway activity in both PDOXs with differential tumor responses consistent with *in vitro* studies. SJ-DIPGX7 showed relative resistance to MEK inhibition and a more moderate impact of combined PI3K/mTOR inhibition compared with the more responsive SJ-DIPGX37, where the drug combination drove greater cytotoxic and cytostatic responses and extended survival of mice carrying intracranial tumors. Pharmacokinetic analyses showed no substantial drug-drug interaction changing drug exposures, so the enhanced effects are likely to represent the result of combined pathway inhibition. However, levels of mirdametinib used to effectively block signaling in our study were significantly higher than those observed in humans following exposure at the current clinical dosing of 4mg twice daily^65^. Given the compelling survival advantage and observed *in vivo* toxicity, there is a rationale to consider local delivery of PI3K/mTOR and MEK inhibitors to these tumors to avoid systemic toxicities and reach the necessary levels of drug activity.

Further studies are needed to understand the intrinsic differences in sensitivity between tumors to ultimately achieve the promise of personalized medicine. Major advances in the treatment of this heterogeneous group of intractable brain tumors will require significant new insights into mechanisms driving disease pathogenesis and even more extensive preclinical testing. Models with a detailed molecular characterization that can be experimentally manipulated and studied *in vitro* and *in vivo* provide powerful tools needed to address this challenge. To facilitate an in-depth exploration of this collection of 21 new PDOX and eight new cell lines, the Pediatric Brain Tumor Portal provides an interactive interface to access and explore all clinical and molecular data and drug screening results including summary overviews for each model (Supplementary Fig. 5) and tools to generate customized mutation oncoprints, gene expression heatmaps, and overlays of dose-response curves for selected drugs and cell lines. The well-characterized models reported here provide a rich resource for the pediatric brain tumor community.

## MATERIALS AND METHODS

### Patient material

Genomic analyses of patient material and use of patient tumor samples to establish xenografts and cell lines were performed with informed consent and approval from the St Jude Institutional Review Board.

### PDOX establishment

Surgical or autopsy tumor samples were transported in neurobasal or DMEM/F12 media without additives at 4°C. Most tissues were processed directly, although samples stored overnight at 4°C also successfully engrafted to form tumors. Tissues were dissociated into a single-cell suspension by gentle pipetting in warm neurobasal media or by enzymatic dissociation with papain as described^66^. Intracranial implantation of 2 × 10^5^ cells - 1 × 10^6^ cells in matrigel into CD-1 nude mice was completed as described^67^. Mice were monitored daily for neurological and health symptoms and euthanized at a humane end point. PDOX tumors were dissected from moribund mice, dissociated, and passaged into 5-10 recipient mice, or cryopreserved in either Millipore or Sigma cell freezing medium (see key resources table in Supplementary Methods). Models were considered established after successfully engrafting through 3 passages. Recovery from cryopreserved cells typically showed delayed *in vivo* tumor growth of 1.5 to 2 times compared to passaging from fresh tumors. Many lines were transduced with a lentivirus to express luciferase and yellow fluorescent protein (CL20-luc2aYFP) for *in vivo* imaging. Mice were maintained in an accredited facility of the Association for Assessment of Laboratory Animal Care in accordance with NIH guidelines. The Institutional Animal Care and Use Committee of SJCRH approved all procedures in this study.

### Cell line propagation and maintenance

To establish pHGG cell lines SJHGGX2c, SJ-HGGX6c, SJ-HGGX42c, SJ-HGGX39c, SJ-DIPGX7c, SJ-DIPGX9c, SJ-DIPGX29c, and SJ-DIPGX37c, fresh PDOX tumors were dissociated as described^66^ and plated in Corning^®^ Ultra-low Attachment plates in media used for neural stem cells and glial progenitor cells consisting of a 1:1 mixture of Neurobasal™ without phenol red (with 2% of B27 without vitamin A and 1% of N2) and ThermoFisher Knock-Out DMEM/F12 (with 2% of Stempro^®^ neural supplement) supplemented with 20ng/ml of human recombinant EGF, 20ng/ml of human recombinant bFGF, 10ng/ml of human recombinant PDGF-AA and -BB, 1% of Glutamax, 1% of sodium pyruvate, 1% of NEAA, 10mM of HEPES, 2μg/ml of heparin and 1x Primocin. In the first passage, the Miltenyi Biotec Mouse Cell Depletion Kit was used, and removal of residual mouse cells was verified by demonstrating successful PCR amplification with human, but not mouse-specific, primer sets for *H3F3A/h3f3a.* Sequences of PCR products were verified. Immunofluorescent staining with anti-human mitochondria and anti-human nuclear antigen antibodies was performed to further verify that cultured cells were of human origin.

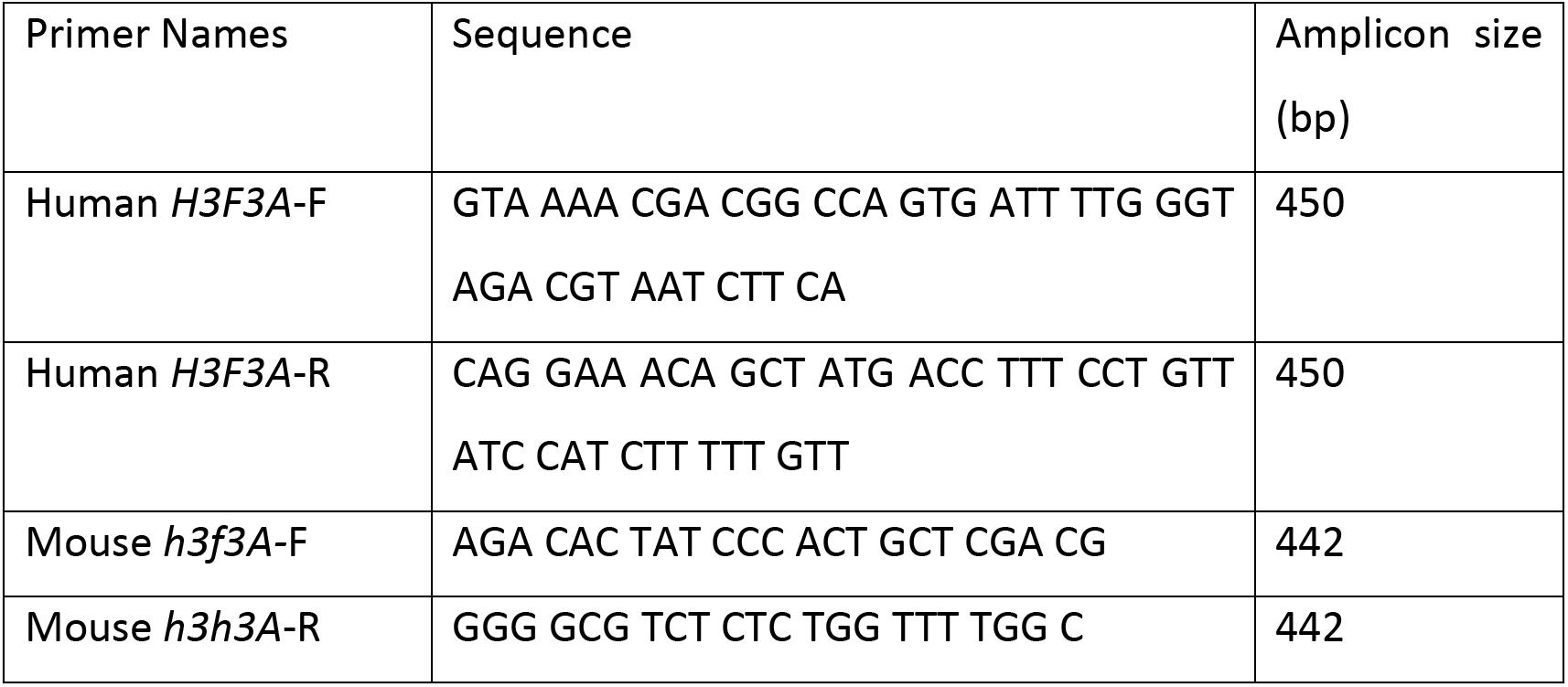

The cell lines were either maintained in suspension culture as tumorspheres or on human ESC-qualified Geltrex artificial extracellular matrix coated tissue culture surface^68^ at 37°C, 5% CO_2_ and 5% of O_2_.

SU-DIPG-IV, SU-DIPG-VI, SU-DIPG-XIII, SU-DIPG-XVII, SUDIPG-XIX, and SU-DIPG-XXI^17^,^22^,^23^ were generous gifts from Dr. Michelle Monje. Normal cell type references were human neural stem cells induced from H9 ES cells (Invitrogen, N7800-100), Human iPSC-derived astrocytes (Tempo Bioscience), and Human brainstem astrocytes (ScienCell Research Laboratories, #1840).

### Short tandem repeat (STR) profiling

Molecular fingerprinting for PDOXs and cell lines was performed with Promega PowerPlex^®^ 16 HS or PowerPlex Fusion^®^ System (Promega Corporation, Madison, WI).

### DNA methylation profiling and copy number analysis

DNA methylation profiles were evaluated using Illumina Infinium Methylation EPIC BeadChip arrays according to the manufacturer’s instructions. Raw IDAT files from pHGG patient tumors, PDOXs, and cell lines as well as a published reference cohort from^13^ were assessed for quality control and pre-processed using the minfi: package in R^69^. Low-quality samples were excluded from the downstream analysis based on mean probe detection with p-value > 0.01. Relevant tumor subgroups were selected from the reference cohort and then subgroups with more than cases were randomly downsampled to be at most 30 cases. Methylation probes were filtered based on the following criteria: detection p > 0.01 in > 50% of the cohort, nonspecificity based on a published list from^70^, and probes residing on sex chromosomes. Methylation arrays then underwent single-sample Noob normalization to derive beta values, with the top 35,060 most variable probes (probe SD > 0.25) selected for downstream analysis. Distances between samples were calculated based on Pearson’s distance and then visualized with the t-SNE algorithm (Rtsne package v.0.11).

Copy-number alteration (CNA) analysis was done using the conumee package (v 1.18.0) with default parameters. Probe intensities were normalized against a reference set of normal brain tissues profiled by MethylationEPIC array (n=34). CNA were then detected as significant positive or negative deviations, which encompassed more than 50% of the chromosomal arm, from the genomic baseline.

### Neuropathology assessment

Standard hematoxylin and eosin histopathologic preparations of 5-μm formalin-fixed and paraffin-embedded tissue sections from patient tumors and derived PDOXs were centrally reviewed by a board-certified neuropathologist specialized in pediatric CNS tumors (J.C.) and blinded to the origin and genetic alterations of the PDOXs.

### Immunohistochemistry and FISH

Immunohistochemistry (IHC) was performed as previously described^71^. For quantification of active Caspase 3 and phospho-histone H3 IHC, Antibodies are listed in the key resources table in Supplementary Methods. Amplification of *PDGFRA* (4q12) and *MYCN* (2p24) was detected by interphase fluorescence *in situ* hybridization in a Clinical Laboratory Improvement Amendments (CLIA)-certified laboratory with probes developed in-house using the following BAC clones: *PDGFRA* (RP11-231C18 + RP11-601I15) with 4p12 control (CTD-2057N12 + CTD-2588A19) and MYCN (RP11-355H10 + RP11-348M12) with 2q35 control (RP11-296A19 + RP11-384O8).

### Whole-genome and whole-exome sequencing analysis

Both whole-genome sequencing (WGS) and whole-exome sequencing (WES) were performed on the majority of the samples with a few samples having only WGS or WES. Paired-end sequencing was conducted on Illumina HiSeq platform with a 100-bp or 125-bp read length or NovaSeq with 150-bp read length. Paired-end reads from WGS and WES were mapped to GRCh37-lite using BWA-aln and quality checked (QC) as previously described^72–74^. For PDOX samples, mapped reads were cleansed of mouse read contamination by XenoCP^75^. For patient tumors and PDOX samples with matched germline samples, somatic mutations including SNVs and Indels were called and classified as previously described^73,76^. Non-silent mutations including missense, nonsense, in-frame insertion, in-frame deletion, frameshift, and splice mutations were reported. Potentially pathogenic germline variants were reported based on these filters: 1) non-synonymous mutations with variant allele frequency (VAF) > 0.2 and coverage > 10x; 2) minor allele frequency (MAF) in general populations < 1e-3 in ExAC^77^; and 3) REVEL score > 0.5^78^, if available, for missense mutations. For patient tumors and PDOX samples without matched germline samples, variants were called and annotated by Bambino and Medal Ceremony^79,80^; variants meeting these criteria were reported as potentially pathogenic variants: 1) any ‘Gold’ variants annotated by Medal Ceremony with alternative allele count > 4; 2) non-Gold but non-silent variants with VAF > 0.3, alternative allele count > 4, and MAF in general populations < 1e-3 in ExAC^77^, 1000 Genomes, and NHLBI. Mutations in pHGG signature gene list (Supplementary Table 2b) were manually reviewed. Oncoprints were created using the online tool ProteinPaint^81^. CNAs were called and annotated by CONSERTING^82^, and focal CNA covering signature genes were manually reviewed.

### RNA sequencing analysis

Paired-end RNA sequencing (RNA-seq) was conducted on Illumina HiSeq platform with 100-bp read length or 125-bp read length. Paired-end reads from RNA-seq were mapped as previously described^83^. For PDOX samples, mapped reads were cleansed of mouse read contamination by XenoCP. Read counts per gene per sample were quantified by HTSeq-Count v 0.11.2 using level and level 2 transcripts of GENCODE v19 annotation^83^. Expression heatmap was plotted based on log-CPM of genes from three expression signatures across the entire cohort, excluding four samples due to different RNA-seq protocol (patient diagnostic and recurrent samples from SJ-HGGX2, patient and PDOX samples from SJ-HGGX6) and four PDOX samples from earlier passages (one PDOX from SJ-DIPGX29, one PDOX from SJ-HGGX39, and two PDOXs from SJ-DIPGX7). Differential expression analyses were carried out on 16 one-to-one matched pairs of PDOXs from the most recent passages and the patient samples from which they were derived using edgeR (v 3.28.0) and limma (v 3.42.0) following the RNAseq123 workflow^84^. Subsequent gene set enrichment analyses were conducted using a custom R script based on hypergeometric test against hallmark gene sets downloaded from MSigDB v 5.2^85^.

Fusion genes were detected using CICERO^86^.

### High throughput screening (HTS)

Growth curves for each cell line were established in 384-well plates (Corning 3707 or 3765) coated with 40 μls of 1% Geltrex matrix with varying cell numbers to determine optimal seeding density for 7-day treatments (Supplementary Table 4). For HTS, all assays were performed with the negative control (DMSO, 0.35%) and the positive control (staurosporine, 19-35 μM) in parallel. The FDA single-point assays were performed at a final concentration of 33 μM [95% CI 16-46 μM]. Dose-response experiments were performed with 10-point, 3-fold serial dilution (19683-fold concentration range), and the mean top concentration tested was 35 μM [95% CI 16-50 μM] (Supplementary Table 5).

The assay plates were loaded into an automated cell culture compatible LiCONiC incubator (37 °C, 5% CO2 and humidified) (LiCONiC US, Woburn, MA) that was integrated into an automated high-throughput screening system (HighRes Biosolutions, Beverly, MA). All compounds were transferred with a pin tool (V&P Scientific, San Diego, CA) and tested in triplicate, and drug exposure time was 7 days except for the fast-growing iAstro cells (shortened to 3 days). At the end of the experiment, the assay plates were equilibrated to room temperature for 20 minutes. Cell viability was measured using CellTiter-Glo^®^ (Promega, Madison, WI). Luminescent signal was read with an EnVision^®^ multimode plate reader (PerkinElmer, Waltham, MA). Screening experiments were processed, and the results visualized using two in-house developed programs: RISE (Robust Investigation of Screening Experiments) and AssayExplorer.

## Data analysis of drug responses

### Raw data processing – log2 RLU dose-response fits

Raw luminescence relative light unit (RLU) values for each compound at each concentration were log2 transformed, normalized to obtain % activity using the following equation: 100x [(mean(negctrl) − compound) / (mean(negctrl) − mean(posctrl))]; and then pooled from replicate experiments prior to fitting. Here, negctrl and posctrl refer to the negative (DMSO) and positive controls (staurosporine) on each plate.

Dose-response curves were fit using the drc^87^ package in R [R Core Team (2012). R: A language and environment for statistical computing. R Foundation for Statistical Computing, Vienna, Austria. ISBN 3-900051-07-0, URL http://www.R-project.org/]. Both a three-parameter (with y0, the response without drug, set to zero) and a four-parameter model (y0 allowed to vary) were fit using the sigmoidal function *LL2.4.* The hill slope was constrained to be between −10 and 0, and EC_50_ was constrained to be between 10^−11^ and 10^−4^ (which roughly equated to the drug concentration range tested in these experiments). For the three-parameter model, yFin, the maximum response of the dose-response curve, was constrained to be between zero and the maximum of the median activities calculated at each concentration overall pooled measurements. For the four-parameter model, y0 and yFin were both constrained to be between the minimum and the maximum of the median activities calculated at each concentration overall pooled measurements. The model with the lowest corrected Akaike information criterion (AICc) was selected as the best fit model.

Area under the curve (AUC) was calculated from the fitted curve using the trapezoid rule in the concentration range 10^−11^ and 10^−4^ Molar. In the event of a failure to fit a sigmoidal dose-response curve, the *smooth.spline* option in R was used to fit a curve that could be used to determine AUC.

### QC metrics for HTS

The z-prime statistic was calculated using the following formula: 1 - ((3xsd(negctrl)) + (3xsd(posctrl))) / abs(mean(negctrl) − mean(posctrl)).

### BRAID model for quantitative synergy analysis

Drug combination experiments were analyzed using the BRAID response surface model^43^. Raw RLU values were processed as described above for the single-agent experiments, except no log2 transformation was applied.

### *In vivo* testing of paxalisib and mirdametinib

Paxalisib (GDC0084) and mirdametinib (PD0325901) were formulated at 1.8 mg/mL or 2.5 mg/mL, respectively, in 1% Methylcellulose and 1% Tween 80 with sonication. The combination was co-formulated and administered as a single gavage.

### Pharmacodynamic assays

SJ-DIPGX37 or SJ-DIPGX7 tumors were implanted in the brain and treated when mice showed decreased activity consistent with large tumors and confirmed visualization of brain tumors by MRI. Mice were dosed by oral gavage, daily for five days with mirdametinib (17mg/kg), paxalisib (12mg/kg), mirdametinib and paxalisib (17 mg/kg and 12 mg/kg, respectively), or vehicle. Two hours after the last dose, mice were perfused with PBS to remove blood, a small piece of tumor was grossly dissected and snap-frozen for Western blot analysis, and the remainder of the tumor and brain was processed for FFPE histology and IHC.

Long-term treatment at these doses caused toxicity manifested as loss of >20% of body weight. To identify doses tolerated for longer treatment needed for survival studies, we treated CD-1 nude mice without tumors in one of seven arms: 1) vehicle; 2) paxalisib at 8 mg/kg; 3) paxalisib at 12 mg/kg; 4) mirdametinib at 14 mg/kg 5) mirdametinib at 17 mg/kg; 6) paxalisib at 8 mg/kg+mirdametinib at 14 mg/kg or 7) paxalisib at 12 mg/kg and mirdametinib at 17 mg/kg. Three mice per arm were treated for five consecutive days, and then the brains were collected 2 hours after the final dose for pharmacodynamic analysis of pathway inhibition (pAKT and pERK) by Western blotting. An additional 3 mice per arm were treated on cycles of 5 days on, two days off for 3 cycles. Body weight and behavioral changes were monitored daily. Paxalisib (8 mg/kg) and mirdametinib (14 mg/kg) alone and in combination were tolerated without loss of >20% body weight, so these doses were selected for a survival study. Western blots for pharmacodynamic studies were performed as previously described^71^ using antibodies listed in the key resources table.

### Survival study

Cryopreserved SJ-DIPGX37 PDOX cells were thawed, and 2.4 × 10^5^ cells in 7.5^l of Matrigel/mouse were implanted into brains of 24 mice as previously described^67^. Bioluminescence imaging (BLI) was monitored weekly, and fifty days after implantation when all mice reached a threshold BLI total flux > 2×10^5^, they were randomized into four treatment groups (6 mice per group) and treated with vehicle, paxalisib (8mg/kg), mirdametinib (14mg/kg) or paxalisib + mirdametinib daily, 5-days on and 2/3-days off. Mice were euthanized when they reached moribund status.

### Pharmacokinetic (PK) study and analysis

The plasma and brain PK of mirdametinib and paxalisib was studied in non-tumor bearing mice to determine potential for a PK drug-drug interaction (DDI). Mice were dosed mirdametinib and paxalisib daily, either alone or in combination, for up to 5 days. Blood samples were obtained from the retro-orbital plexus under anesthesia or by cardiac puncture upon termination. Brains were harvested after cardiac puncture and aortic perfusion with PBS. Samples were stored at −80 °C until analysis with qualified LC-MS/MS methods. Plasma concentration-time (Ct) data were analyzed using nonlinear mixed effects modeling implemented in Monolix 2019R2 (Lixoft, Antony, France). To enhance modeling precision and power, additional paxalisib plasma Ct data from a separate DDI study with another targeted compound was added to the analysis. Combination status was tested as a covariate upon the apparent clearance (CL/F) of each compound using the likelihood ratio test. Brain concentrations were log transformed and analyzed using two-way ANOVA with sample time and combination status as factors. Practically impactful interactions were defined as a ≥ twofold change in the parameters of interest, consistent with the conventionally accepted threshold for preclinical and DDI studies^44,45^. Additional details are presented in the Supplementary Materials.

## Supporting information

He et al. Supplementary Figures and Methods bioarchive

## Data availability

DNA methylation profiles are available in Gene Expression Omnibus (GEO), accession GSE152035. All whole-genome, whole-exome and RNA-sequencing will be deposited at the European Genome-phenome Archive (EGA), which is hosted by the European Bioinformatics Institute (EBI). Interactive visualizations of data can be explored in the Pediatric Brain Tumor Portal (pbtp.stjude.cloud). Source data are provided with this paper.

## Acknowledgments

We are grateful to the patients and families who donated tissue to support pHGG research. This work was funded in part by the National Brain Tumor Society Defeat Pediatric Brain Tumors Research Collaborative (SJB, CT, AAS), NIH CA096832 (SJB, MFR, JZ), the NCI Cancer Center Support Grant CA21765, Musicians Against Childhood Cancer and ALSAC. We thank the St. Jude Children’s Research Hospital (SJCRH) Research Information Services and Cloud Applications groups for the development of the online Pediatric Brain Tumor Portal, Biomedical Communications for assistance with artwork, Center for In Vivo Imaging and Therapeutics (CIVIT), Hartwell Center, and Biorepository. We thank Drs. Jie Zhang and Ali G. Saad (Le Bonheur Children’s Hospital) for fresh tissue collection, Asli Goktug (SJCRH) for assistance with HTS, Brittney Gordon in the SJCRH DNB Xenograft Core for assistance with PDOX cryopreservation, and Dr. Arzu Onar-Thomas for assistance with statistical analyses.

