## Supplementary material for "Patient-Derived Orthotopic Xenografts and Cell Lines from Pediatric High-Grade Glioma Recapitulate the Heterogeneity of Histopathology, Molecular Signatures, and Drug Response": He et al. Supplementary Figures and Methods bioarchive

**Supplementary Figure 1**: **Heatmap of large-scale copy number changes in patient tumors, PDOX, and cell lines.** Patient tumor and matched PDOX and cell lines are grouped together in columns. Abbreviated line numbers are listed across the top. Rows show large scale gain (red) or loss (blue) of chromosome arms indicated at right. Scored large-scale copy number changes encompassed more than 50% of a chromosome arm.

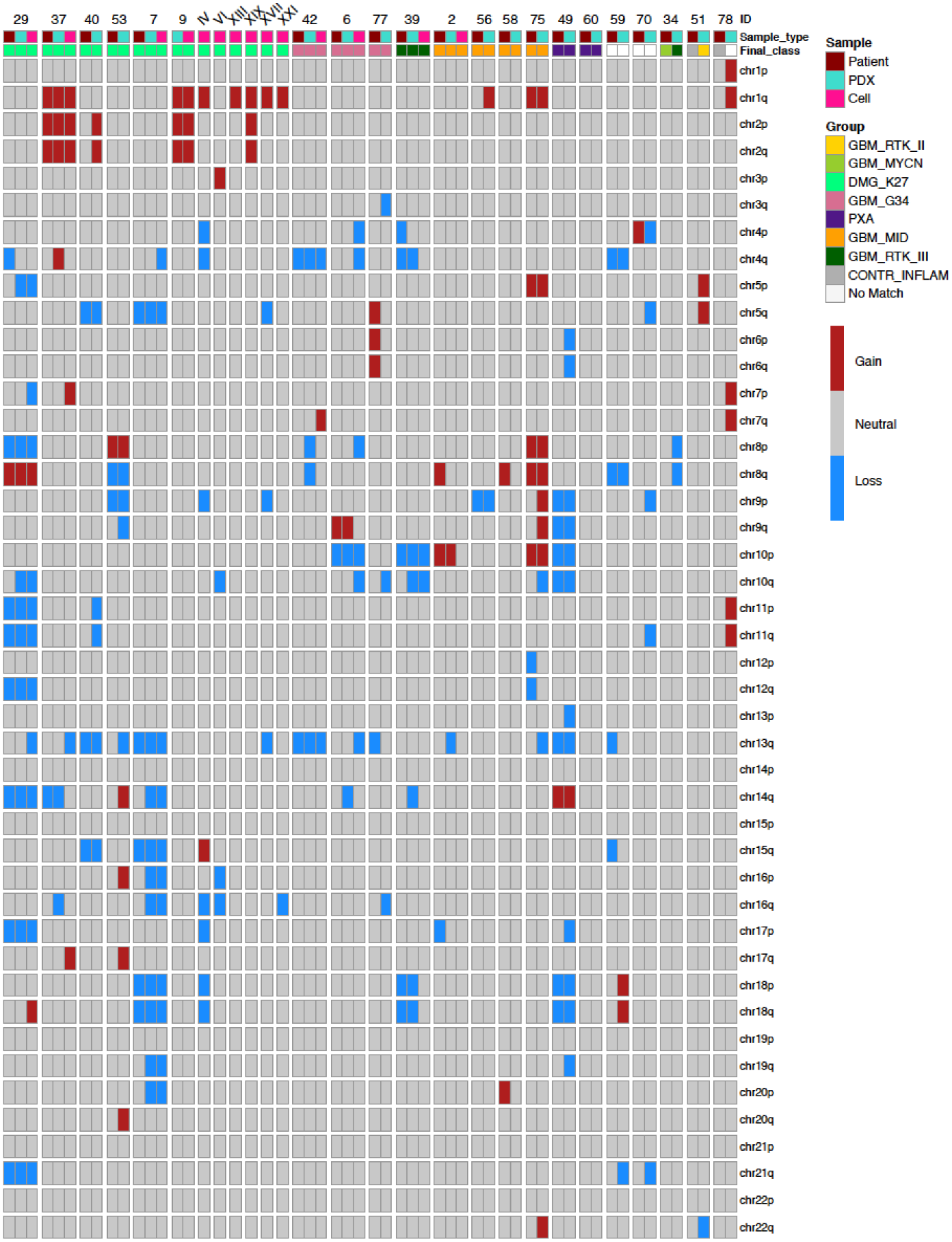

**Supplementary Figure 2**: **Extra-chromosomal DNA amplifications in PDOX models.** (a) WGS (top 3 tracks) and RNA-seq (lower 3 tracks) showed amplification and overexpression, respectively, of a TMEM165-PDGFRA fusion gene in SJ-HGGX75 patient tumor and matched PDOX compared to normal copy number and expression in SJ-HGGX70. The structural variant connecting the boundaries of the amplified segment was identified in WGS of SJ-HGGX75 patient tumor and matched PDOX. (b) TMEM165-PDGFRA fusion gene was identified from RNA-seq data using CICERO in SJ-HGGX75 patient tumor and matched PDOX. The resulting chimeric protein replaces the most N-terminal Ig-like domains of PDGFR$\alpha$ with 70 amino acids from the N-terminus of TMEM165. (c) Interphase fluorescence *in situ* hybridization of PDOX SJ-HGGX75 shows amplification of *PDGFRA* (red, 4q12; control, green, 4p12) in the form of double minutes in 100% of the 200 evaluable nuclei. (d) WGS (top 3 tracks) and RNA-seq (lower 2 tracks) showed *MYCN* amplification and overexpression, respectively, in the PDOX from SJ-DMGX40, but not the diagnostic tumor from which it was derived, or autopsy from the same patient. (e) Interphase fluorescence *in situ* hybridization of PDOX SJ-DMGX40 shows amplification of *MYCN* (red, 2p24; control, green, 2q35) in the form of double minutes in 100% of the 200 evaluable nuclei. Scale bar in c,e: 15 μm

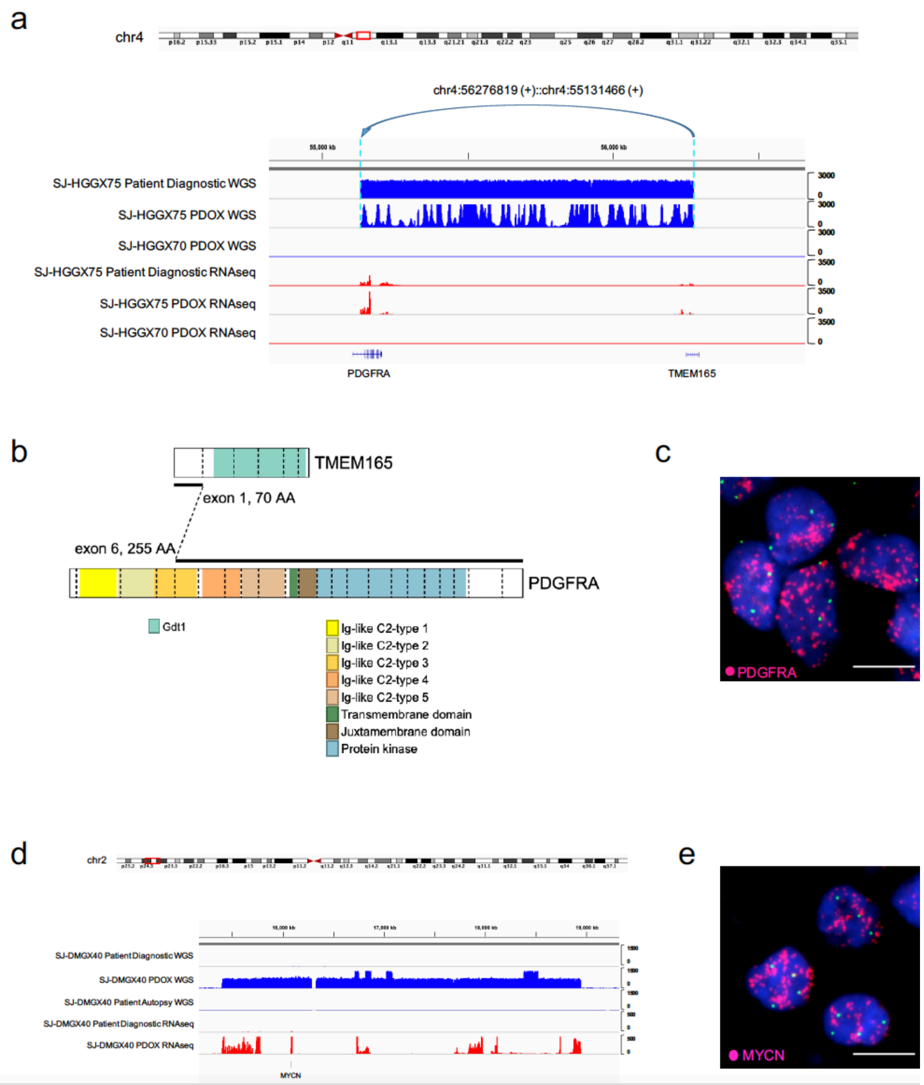

**Supplementary Figure 3: Expression signatures of PDOX models recapitulate primary tumors from which they are derived.** Pearson correlation coefficient of RNA-seq quantification (logCPM) between each PDOX sample and all 16 patient tumors that have matched PDOXs. The red dots represent the Pearson correlation coefficient between each PDOX sample and the matched patient tumor, black dots show correlation between each PDOX sample and other patient tumors. Genes include all mRNA genes except those identified as genes differentially expressed between all PDOX compared to patient tumors (Supplementary Table 3).

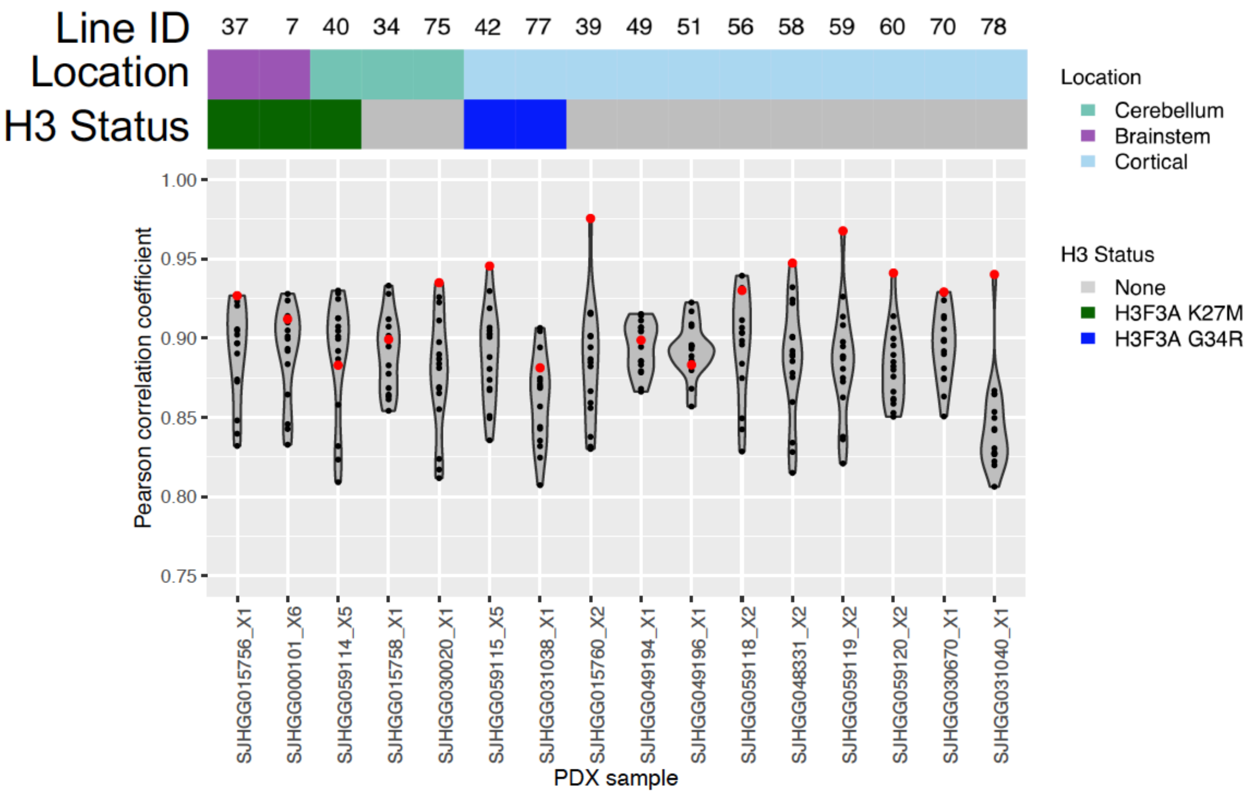

**Supplementary Figure 4. Fidelity of expression signatures in PDOX models from different passages, and in cell lines compared with the PDOX from which they were derived.** Scatterplot comparing expression in indicated PDOX and cell lines (RNAseq, $log_2$-counts per million).

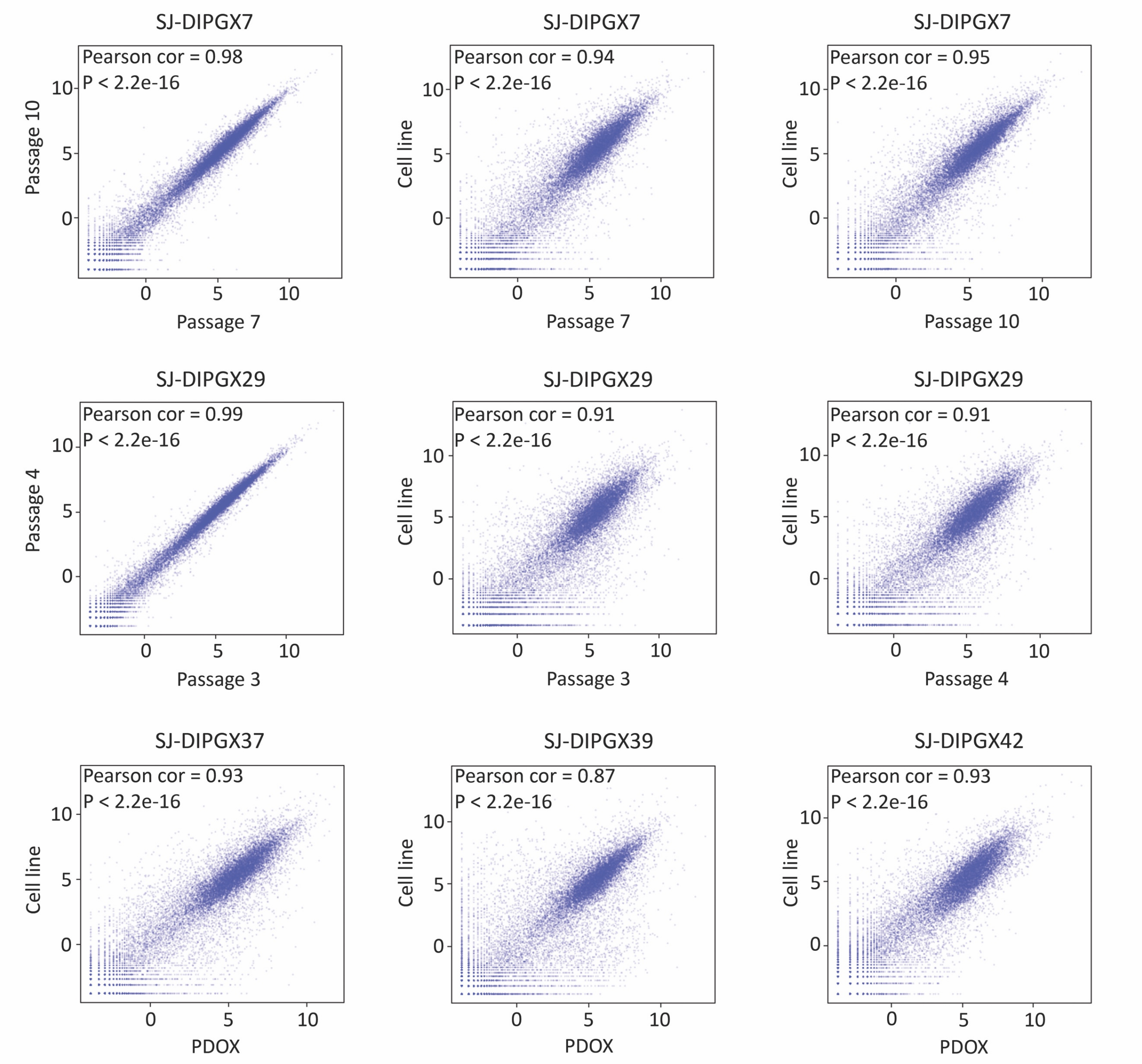

**Supplementary Figure 5: Characterization sheet for pHGG models available through Pediatric Brain Tumor Portal.** Characterization sheet for PDOX line SJ-HGGX42 and associated samples is shown here as a representative example. PDF summaries for each model can be accessed and downloaded from the PBTP, and this information and multiple additional options for interactive exploration of data from this paper are available through the online portal.

### Pediatric Brain Tumor Portal

#### Summary

**Name:** SJ-HGGX42
**Sample ID:** SJHGG059115
**Disease:** High-grade Glioma
**Associated Acronyms:** HGG, GBM
**Histone H3 Status:** H3.3 G34R

#### Clinical Information

**Clinical Group:** Cortical HGG
**Pathologic Group:** Glioblastoma
**Grade:** IV
**Patient Sample Type:** Diagnostic
**Age at Diagnosis:** 13 years
**Survival from Diagnosis:** 12 months
**Sex:** Male
**Patient Tumor Location:** Cortical
**Treatment History:** Treatment Naive

#### Sample Information

**PDOX Label:** YFP-Luciferase
**Cell Line Label:** YFP-IRES-Luciferase
**Mouse Survival (PDOX→PDOX):** 5 months
**Mouse Survival (cell culture→PDOX):** 5 months
**Material Available:** PDOX, Cell line

#### Sequence Source and Oncoprint

| Sample | Sample ID | WGS | WES | RNAseq | Methylation |
|---|---|---|---|---|---|
| Patient Germline | SJHGG059115_G1 | N/A | N/A | N/A | N/A |
| Patient Tumor (Diagnostic) | SJHGG059115_D1 | ✓ | ✓ | ✓ | ✓ |
| PDOX Tumor (Derived from Diagnostic) | SJHGG059115_X5 | ✓ | ✓ | ✓ | ✓ |
| Cell line (Derived from PDOX) | SJHGG054830_C2 | N/A | ✓ | ✓ | ✓ |
| PDOX Tumor (Derived from cell line) | SJHGG059115_X31 | ✓ | ✓ | ✓ | ✓ |
| Patient Tumor (Autopsy) | SJHGG059115_A1 | ✓ | ✓ | ✓ | ✓ |

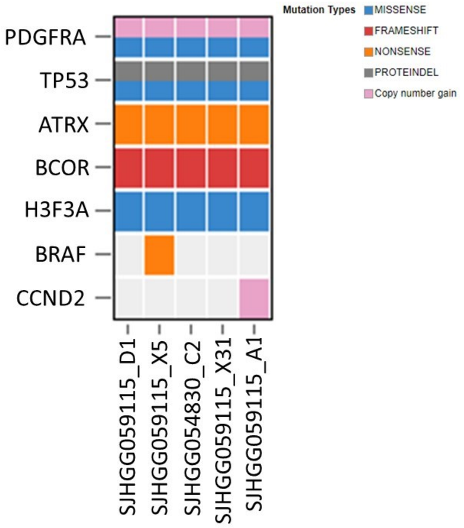


**Table** indicates sequencing performed (✓); WGS, whole genome sequencing; WES, whole exome sequencing; RNAseq, RNA sequencing; Methylation, BeadChip array profiling human DNA methylation; N/A, sequence not available.

**Oncoprint** for pediatric high-grade glioma signature mutations in these samples.

#### Histology

### H&E

Patient Tumor

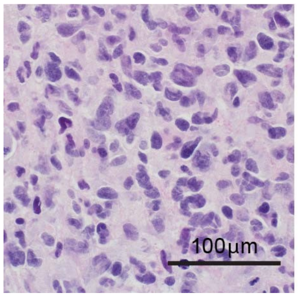


PDOX Tumor

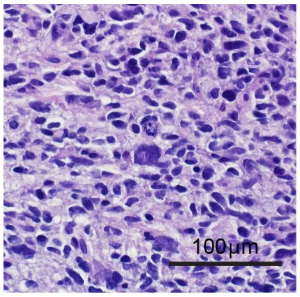


##### ATRX

Patient Tumor

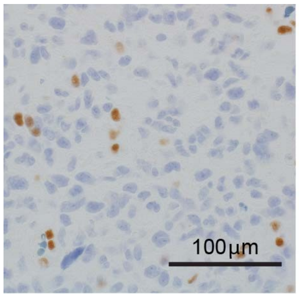


PDOX Tumor

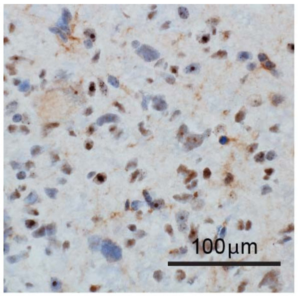

#### Mutation Status

##### ATRX

| Sample | aa change |
|---|---|
| SJHGG059115_D1 | S692* |
| SJHGG059115_X5 | S692* |
| SJHGG054830_C2 | S692* |
| SJHGG059115_X31 | S692* |
| SJHGG059115_A1 | S692* |

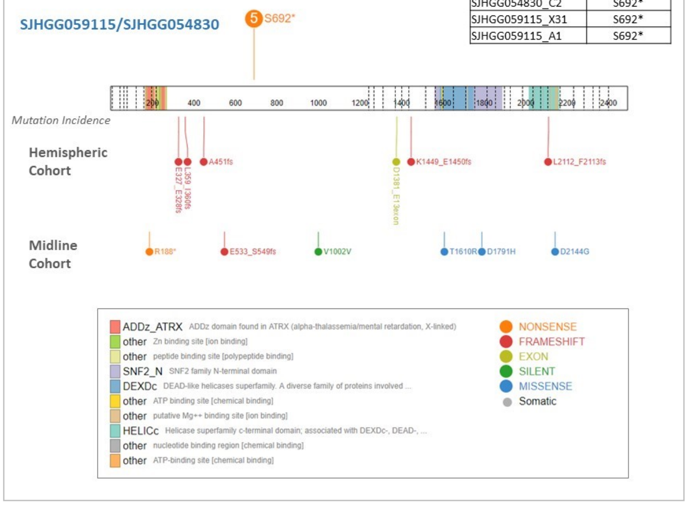


##### BCOR

| Sample | aa change |
|---|---|
| SJHGG059115_D1 | S177fs |
| SJHGG059115_X5 | S177fs |
| SJHGG054830_C2 | S177fs |
| SJHGG059115_X5 | S177fs |
| SJHGG059115_X5 | S177fs |

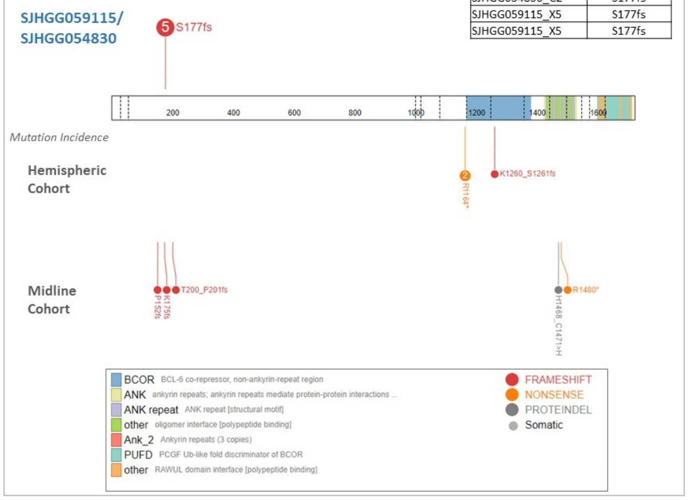


##### BRAF

| Sample | aa change |
|---|---|
| SJHGG059115_D1 | -- |
| SJHGG059115_X5 | S316* |
| SJHGG054830_C2 | -- |
| SJHGG059115_X31 | -- |
| SJHGG059115_A1 | -- |

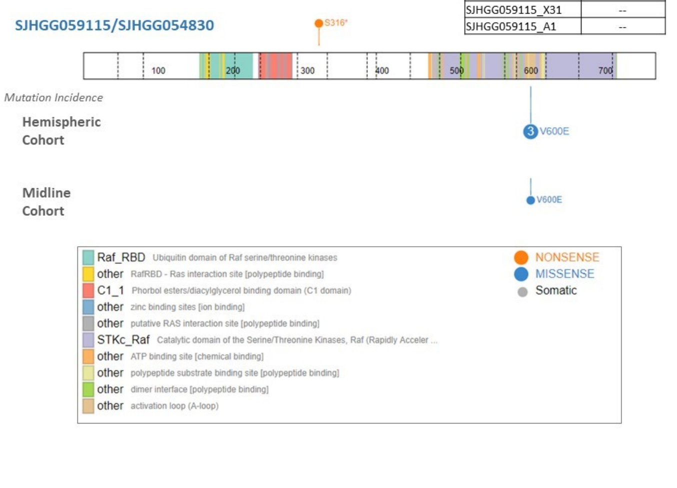


##### PDGFRA

| Sample | aa change |
|---|---|
| SJHGG059115_D1 | P343S |
| SJHGG059115_X5 | P343S |
| SJHGG054830_C2 | P343S |
| SJHGG059115_X31 | P343S |
| SJHGG059115_A1 | P343S |

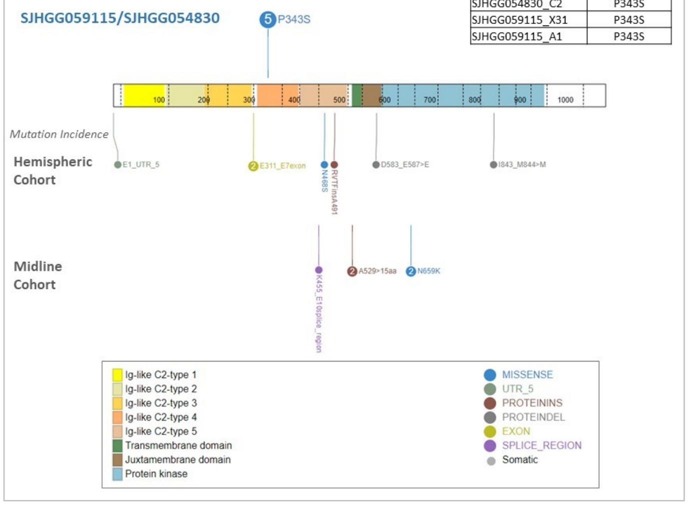


### TP53

| Sample | aa change |
|---|---|
| SJHGG059115_D1 | L264_L265>L |
| SJHGG059115_D1 | R248Q |
| SJHGG059115_X5 | L264_L265>L |
| SJHGG059115_X5 | R248Q |
| SJHGG054830_C2 | L264_L265>L |
| SJHGG054830_C2 | R248Q |
| SJHGG059115_X31 | L264_L265>L |
| SJHGG059115_X31 | R248Q |
| SJHGG059115_A1 | L264_L265>L |
| SJHGG059115_A1 | R248Q |

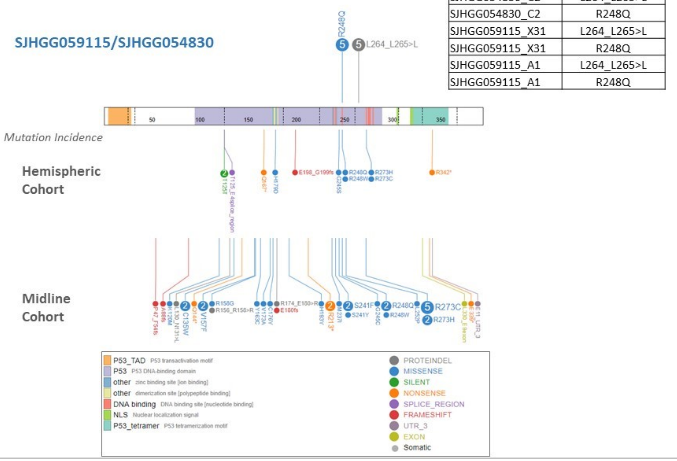


### H3F3A

| Sample | aa change |
|---|---|
| SJHGG059115_D1 | G34R |
| SJHGG059115_X5 | G34R |
| SJHGG054830_C2 | G34R |
| SJHGG059115_X31 | G34R |
| SJHGG059115_A1 | G34R |

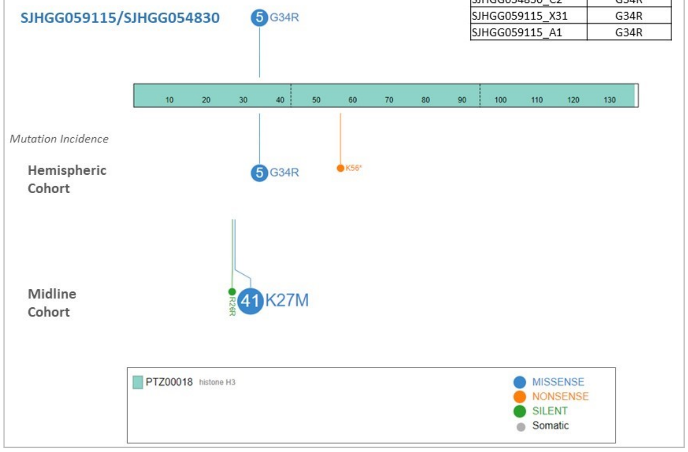


Figure legend, see page 3

Mutation Status figure legend:

Protein paint diagrams show schematic of protein with functional domains shown in legend box. Mutations associated with samples for this line are shown above the protein diagram. Mutations in cohort of 127 pediatric HGGs separated as hemispheric and midline tumors from [1] are shown below. Number of samples with the specific mutation are listed in the circle marking the mutated residue.

#### Copy Number Variation

Patient Tumor
SJHGG059115_D

PDOX Tumor
SJHGG059115_X

Cell line*
SJHGG054830_C

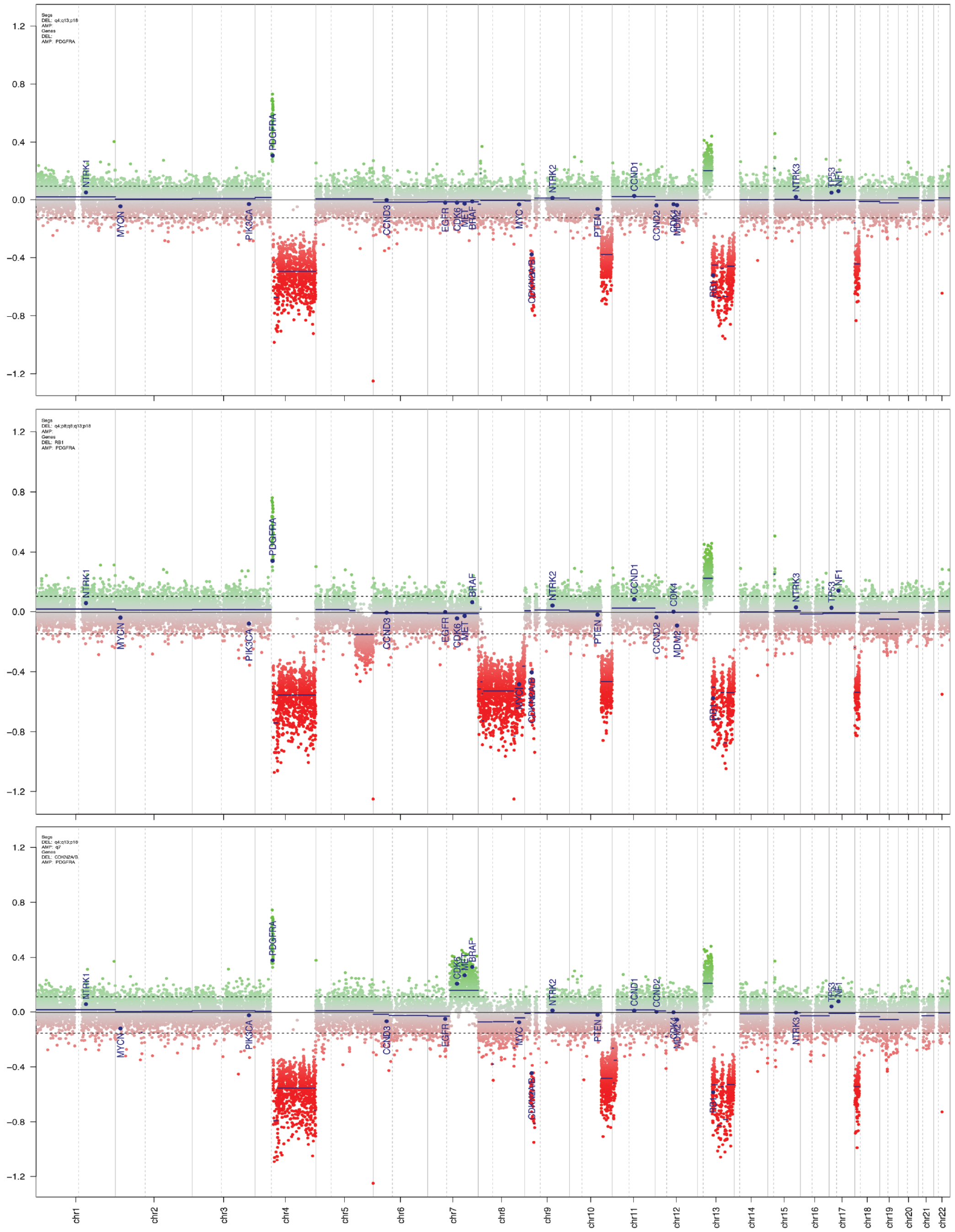


*Cell line derived from PDOX

DNA copy number variation analysis was performed from methylation array data using Conumee [2]. The Y axis shows the log2 copy number ratio of the tumor sample compared to a panel of normal reference brain tissues. Copy number ratios are plotted across chromosomes with the dotted vertical lines representing centromeres. Chromosomal gains or losses are detected as significant positive or negative deviations from genomic baseline. Brain tumor relevant gene regions are highlighted for easier assessment of chromosomal or focal alterations.

#### Brain Tumor Methylation Classification

t-SNE plot showing glioma subgroups based on DNA methylation profiling. Patient tumors (circles), PDOX (squares) and cell lines (diamonds) reported here are outlined in black and reference samples from [3] shown without black outline. Enlarged area shows the methylation class cluster for this sample, and circled samples show the matched samples for this PDOX.

| Sample | Methylation Class |
| --- | --- |
| Patient Tumor | glioblastoma, IDH wildtype, H3.3 G34 mutant |
| PDOX Tumor | glioblastoma, IDH wildtype, H3.3 G34 mutant |
| Cell Line (derived from PDOX) | glioblastoma, IDH wildtype, H3.3 G34 mutant |

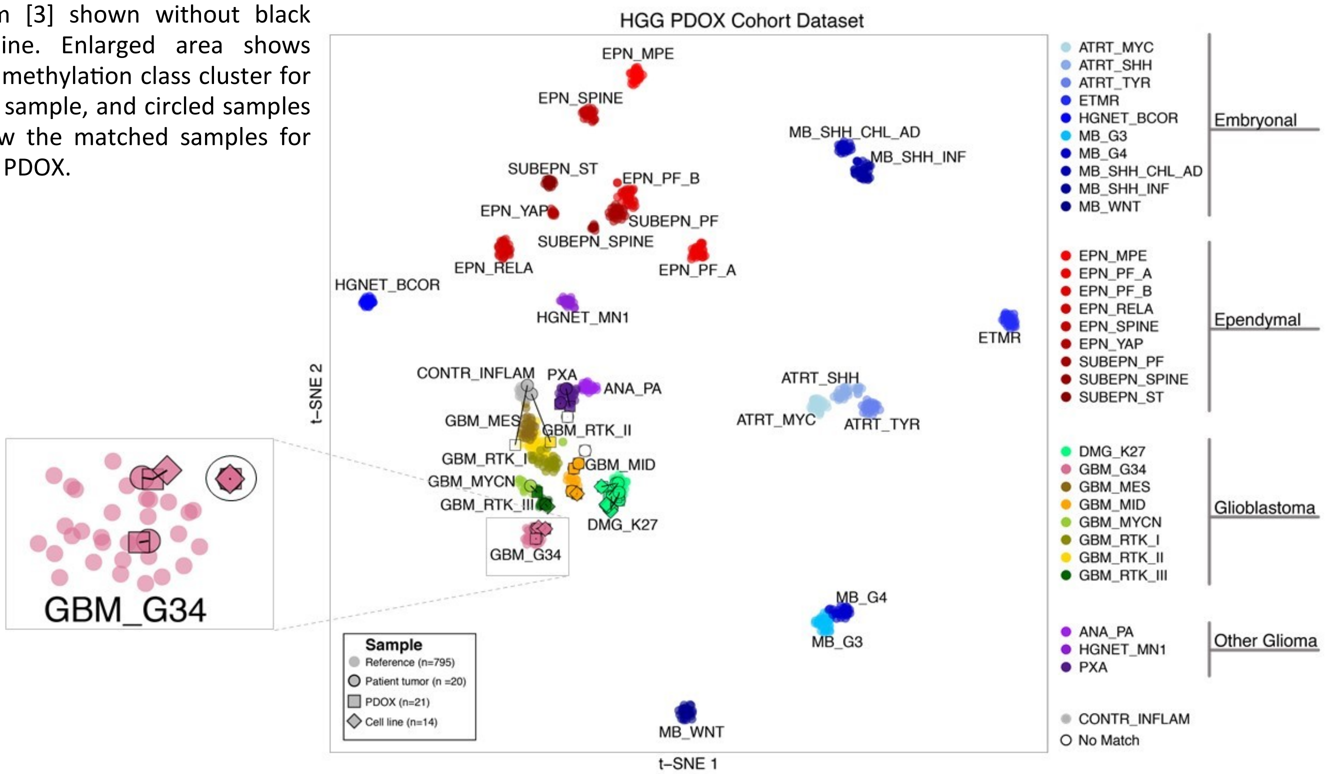


#### Short Tandem Repeat (STR) DNA Typing

| Marker | Allele 1 | Allele 2 |
| --- | --- | --- |
| Amelogenin | X | Y |
| CSF1PO | 10 | 11 |
| D13S317 | 13 | |
| D16S539 | 9 | 11 |
| D18S51 | 12 | 18 |
| D21S11 | 29 | 30 |
| D3S1358 | 14 | 15 |
| D5S818 | 13 | |
| D7S820 | 11 | 12 |
| D8S1179 | 13 | |
| THO1 | 6 | 9.3 |
| TPOX | 9 | 11 |
| vWA | 17 | 18 |

DNA fingerprint from Promega PowerPlex 16 STR assay.


When publishing or referencing data, please cite the St. Jude Children's Research Hospital Pediatric Brain Tumor Portal and include the URL: **https://pbtp.stjude.cloud**

#### Resource

The Pediatric Brain Tumor Portal, organized by the St. Jude Neurobiology and Brain Tumor Program, shares resources and data to support basic and translational investigation of pediatric brain tumors. We look forward to your questions as we continue to provide this resource for the research community.

Please contact us at ****

**Supplementary Figure 6: DIPG cells grown as tumorspheres or adherent cultures on Geltrex basement membrane matrix respond similarly when tested with 53 drugs representing a range of different mechanisms of action.**

Scatterplot of AUCs testing dose-response for 53 compounds in SJ-DIPGX7c grown under adherent vs neurosphere conditions. The Pearson correlation is 0.994. Black line has slope = 1.

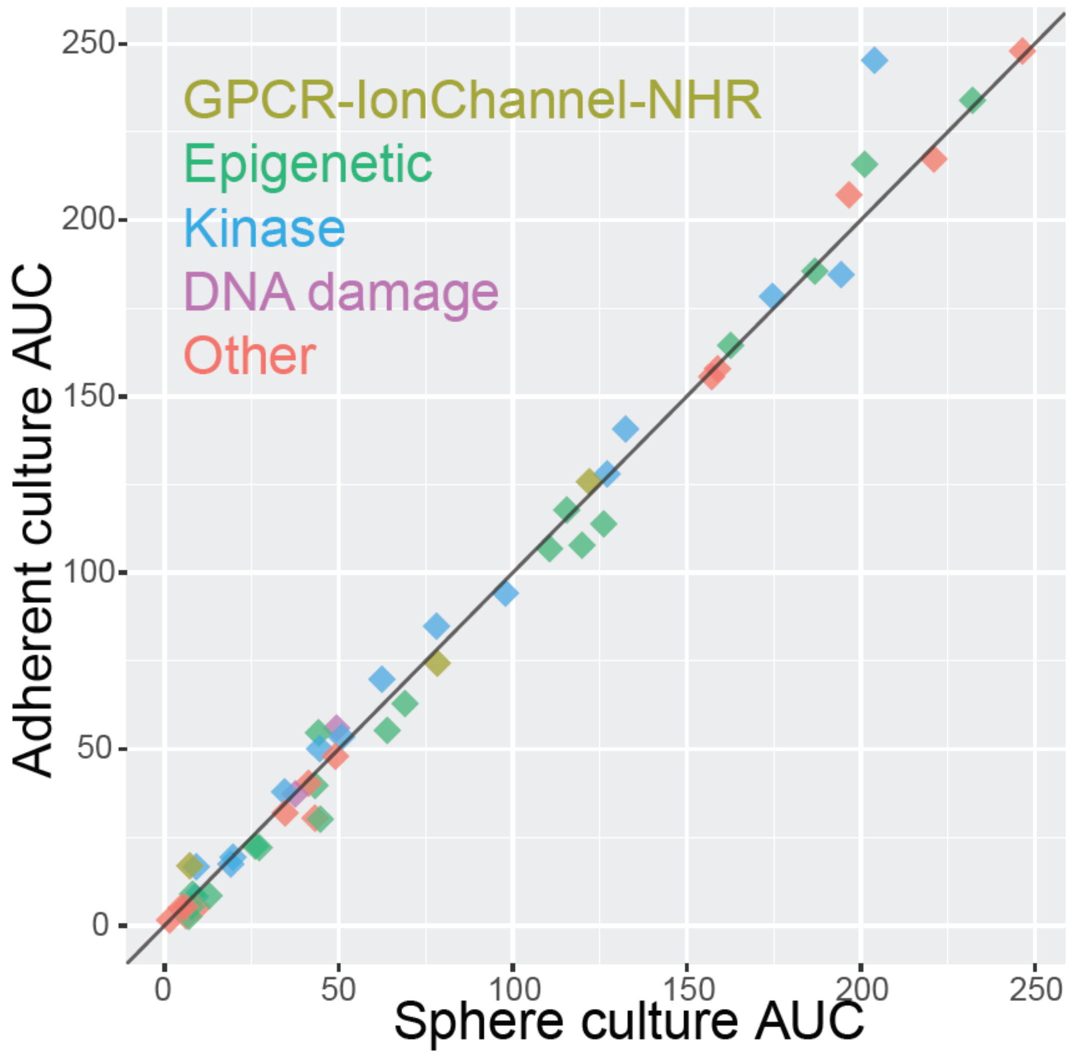

**Supplementary Figure 7: Z prime values for drug screening in 16 cell lines from 310 384-well assay plates.** Scatterplot of Z prime values for each plate screened in this study color coded by cell lines assayed. A Z prime score between 0.5 and 1 indicates an excellent signal to noise ratio.

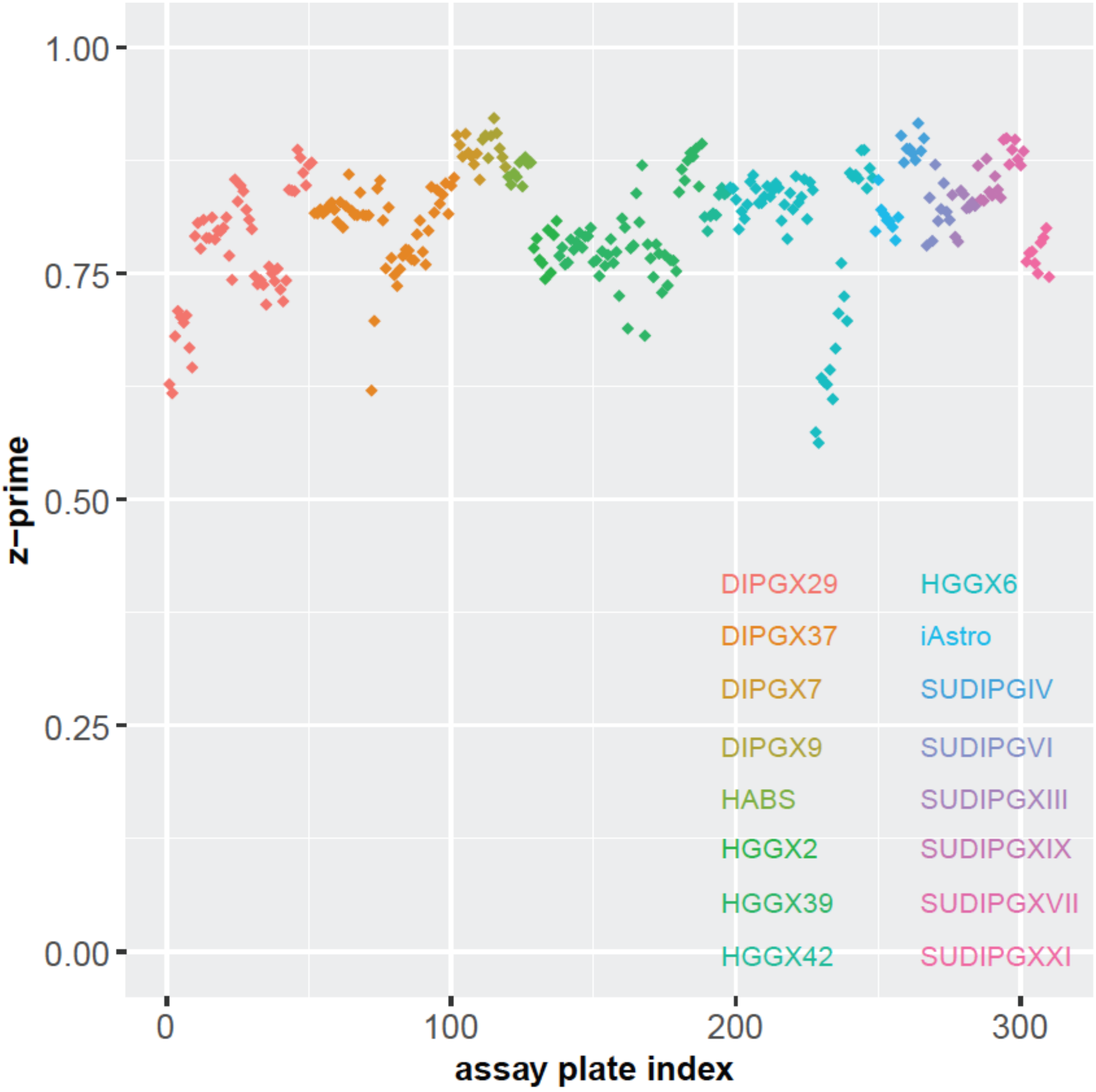

**Supplementary Figure 8: Additional analysis of the screening results from 93 compounds across 14 pHGG models and two normal astrocyte lines.** (a) Select dose-response curves for the drugs highlighted in Figure 6b. iAstro is depicted in black dashed lines, HABS in black solid lines, and pHGG models are colored gray or by histone mutation status where DIPGX37 is indicated. (b) Unsupervised hierarchical clustering of drug AUC z-scores for all compounds tested. Column and column labels are color coded by histone mutation status. Each row represents a single compound and is annotated by mechanism of action with color code shown above. The color code for histone mutation status is: H3-wt (red), H3.3 G34R (blue), H3.1 K27M (turquoise), and H3.3 K27M (green). Control cell lines (iAstro and HABS) are black.

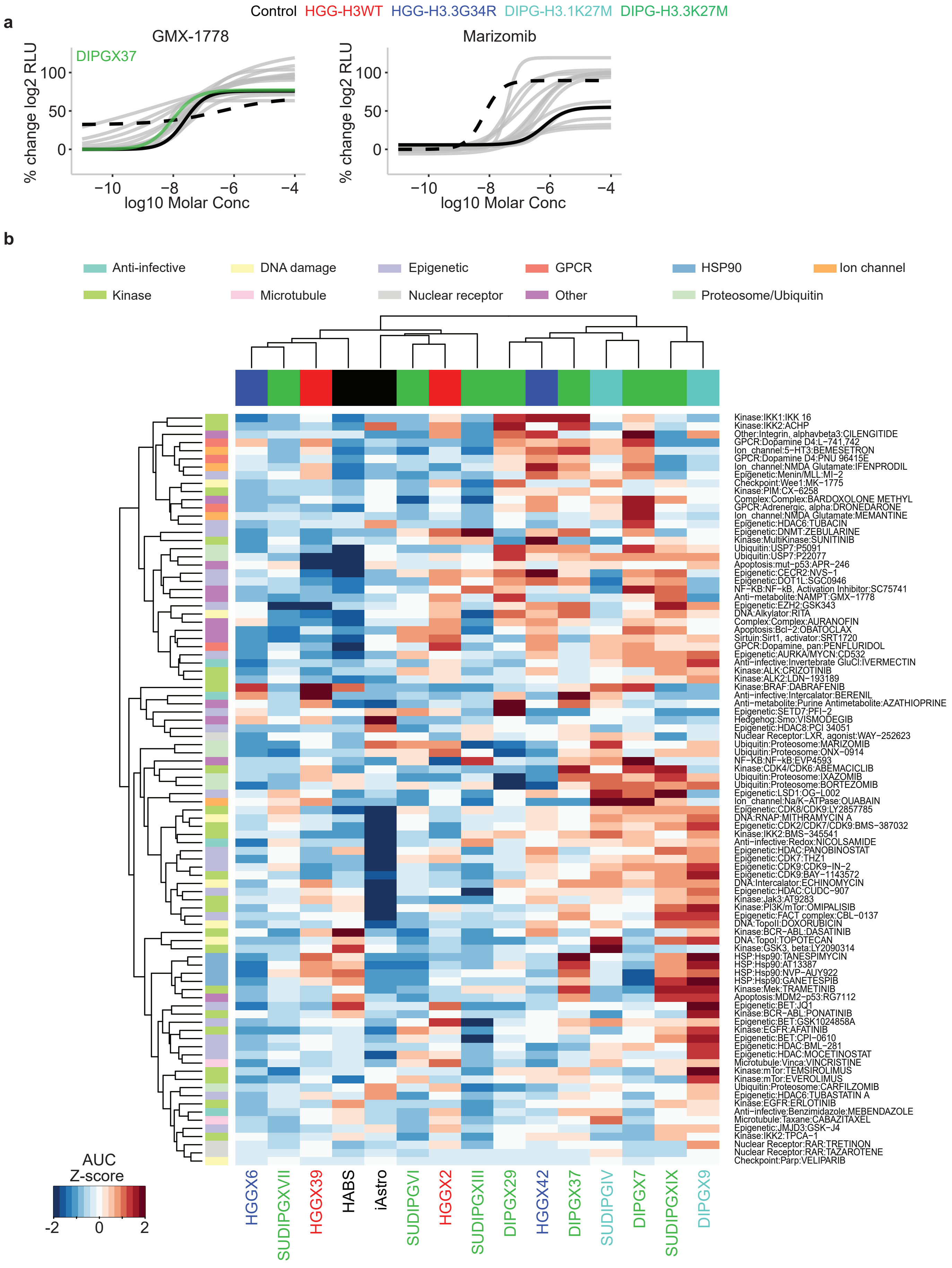
a
Control HGG-H3WT HGG-H3.3G34R DIPG-H3.1K27M DIPG-H3.3K27M
GMX-1778
DIPGX37
Marizomib
% change log2 RLU
log10 Molar Conc
b
Anti-infective
DNA damage
Epigenetic
GPCR
HSP90
Ion channel
Kinase
Microtubule
Nuclear receptor
Other
Proteosome/Ubiquitin
Kinase:IKK1:IKK 16
Kinase:IKK2:ACHP
Other:Integrin, alphavbeta3:CILENGITIDE
GPCR:Dopamine D4:L−741,742
Ion_channel:5−HT3:BEMESETRON
GPCR:Dopamine D4:PNU 96415E
Ion_channel:NMDA Glutamate:IFENPRODIL
Epigenetic:Menin/MLL:MI−2
Checkpoint:Wee1:MK−1775
Kinase:PIM:CX−6258
Complex:Complex:BARDOXOLONE METHYL
GPCR:Adrenergic, alpha:DRONEDARONE
Ion_channel:NMDA Glutamate:MEMANTINE
Epigenetic:HDAC6:TUBACIN
Epigenetic:DNMT:ZEBULARINE
Kinase:MultiKinase:SUNITINIB
Ubiquitin:USP7:P5091
Ubiquitin:USP7:P22077
Apoptosis:mut−p53:APR−246
Epigenetic:CECR2:NVS−1
Epigenetic:DOT1L:SGC0946
NF−KB:NF−kB, Activation Inhibitor:SC75741
Anti−metabolite:NAMPT:GMX−1778
Epigenetic:EZH2:GSK343
DNA:Alkylator:RITA
Complex:Complex:AURANOFIN
Apoptosis:Bcl−2:OBATOCLAX
Sirtuin:Sirt1, activator:SRT1720
GPCR:Dopamine, pan:PENFLURIDOL
Epigenetic:AURKA/MYCN:CD532
Anti−infective:Invertebrate GluCl:IVERMECTIN
Kinase:ALK:CRIZOTINIB
Kinase:ALK2:LDN−193189
Kinase:BRAF:DABRAFENIB
Anti−infective:Intercalator:BERENIL
Anti−metabolite:Purine Antimetabolite:AZATHIOPRINE
Epigenetic:SETD7:PFI−2
Hedgehog:Smo:VISMODEGIB
Epigenetic:HDAC8:PCI 34051
Nuclear Receptor:LXR, agonist:WAY−252623
Ubiquitin:Proteosome:MARIZOMIB
Ubiquitin:Proteosome:ONX−0914
NF−KB:NF−kB:EVP4593
Kinase:CDK4/CDK6:ABEMACICLIB
Ubiquitin:Proteosome:IXAZOMIB
Ubiquitin:Proteosome:BORTEZOMIB
Epigenetic:LSD1:OG−L002
Ion_channel:Na/K−ATPase:OUABAIN
Epigenetic:CDK8/CDK19:LY2857785
DNA:RNAP:MITHRAMYCIN A
Epigenetic:CDK2/CDK7/CDK9:BMS−387032
Kinase:IKK2:BMS−345541
Anti−infective:Redox:NICOLSAMIDE
Epigenetic:HDAC:PANOBINOSTAT
Epigenetic:CDK7:THZ1
Epigenetic:CDK9:CDK9−IN−2
Epigenetic:CDK9:BAY−1143572
DNA:Intercalator:ECHINOMYCIN
Epigenetic:HDAC:CUDC−907
Kinase:Jak3:AT9283
Kinase:PI3K/mTor:OMIPALISIB
Epigenetic:FACT complex:CBL−0137
DNA:TopoII:DOXORUBICIN
Kinase:BCR−ABL:DASATINIB
DNA:TopoI:TOPOTECAN
Kinase:GSK3, beta:LY2090314
HSP:Hsp90:TANESPIMYCIN
HSP:Hsp90:AT13387
HSP:Hsp90:NVP−AUY922
HSP:Hsp90:GANETESPIB
Kinase:Mek:TRAMETINIB
Apoptosis:MDM2−p53:RG7112
Epigenetic:BET:JQ1
Kinase:BCR−ABL:PONATINIB
Epigenetic:BET:GSK1024858A
Kinase:EGFR:AFATINIB
Epigenetic:BET:CPI−0610
Epigenetic:HDAC:BML−281
Epigenetic:HDAC:MOCETINOSTAT
Microtubule:Vinca:VINCRISTINE
Kinase:mTor:TEMSIROLIMUS
Kinase:mTor:EVEROLIMUS
Ubiquitin:Proteosome:CARFILZOMIB
Epigenetic:HDAC6:TUBASTATIN A
Kinase:EGFR:ERLOTINIB
Anti−infective:Benzimidazole:MEBENDAZOLE
Microtubule:Taxane:CABAZITAXEL
Epigenetic:JMJD3:GSK−J4
Kinase:IKK2:TPCA−1
Nuclear Receptor:RAR:TRETINON
Nuclear Receptor:RAR:TAZAROTENE
Checkpoint:Parp:VELIPARIB
AUC
Z-score
HGGX6
SUDIPGXVII
HGGX39
HABS
iAstro
SUDIPGVI
HGGX2
SUDIPGXIII
DIPGX29
HGGX42
DIPGX37
SUDIPGIV
DIPGX7
SUDIPGXIX
DIPGX9

**Supplementary Fig. 9: Pharmacodynamic analyses of mirdametinib and paxalisib show effective MEK and PI3K pathway inhibition in the brain**

Western blots with the indicated antibodies to analyze lysates from: (a) SJ-DIPGX37 intracranial tumors in mice dosed with vehicle (veh, lanes 1-3), 25 mg/kg mirdametinib (mir, lanes 4-6), or 18 mg/kg paxalisib (pax, lanes 7-9) daily for 5 days, and tissue collected 2 hours after last dose. Antibodies are shown at the right. (b) Brain from CD1-nude mice treated with vehicle (lanes 1-3), 8 mg/kg pax (lanes 4-6), 12 mg/kg pax (lanes 7-9), 8 mg/kg pax + 14 mg/kg mir (lanes 10-12) and 12 mg/kg pax + 17 mg/kg mir (lanes 13-15). (c) Brain from CD1-nude mice treated with vehicle (lanes 1-3), 14 mg/kg mir (lanes 4-6), 17 mg/kg mir (lanes 7-9), 14 mg/kg mir + 8 mg/kg pax (lanes 10-12) and 17 mg/kg mir + 12 mg/kg pax (lanes 13-15).

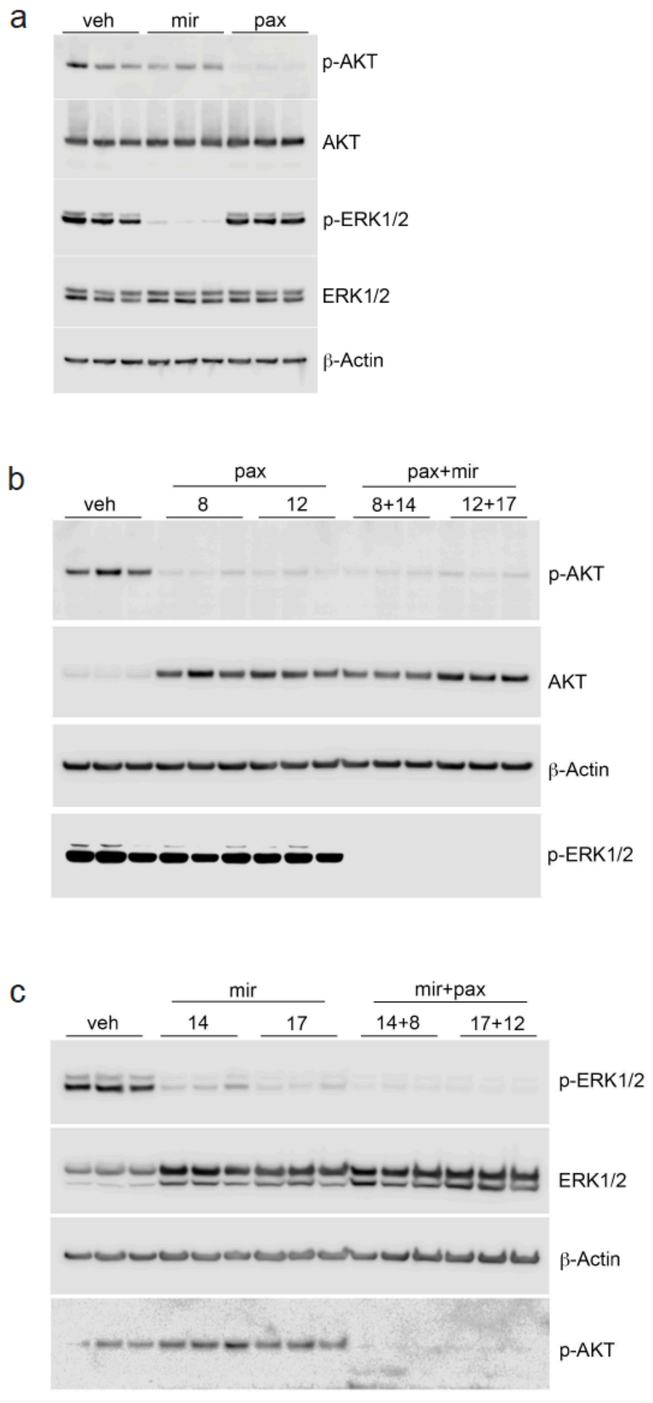

**Supplementary Fig 10: Pharmacokinetic (PK) and pharmacodynamic analyses of Paxalisib and Mirdametinib show lack of appreciable plasma or brain PK drug interaction, high brain exposure, and effective pathway inhibition**

a. Mirdametinib population mean and 90% prediction interval plasma concentration-time profiles alone (mir) and in combination with Paxalisib (mir+pax). A minor increase in Mirdametinib AUC (1.30-fold) was observed in combination.
b. Paxalisib population mean and 90% prediction interval plasma concentration-time profiles alone (pax) and in combination with Mirdametinib (pax+mir). A minor increase in Paxalisib AUC (1.63-fold) was observed in combination.
c. Mirdametinib mean and standard deviation brain concentration-time profiles alone (mir) and in combination with Paxalisib (mir+pax). Only the 2-hour time point is significantly higher for the combination (907 vs 498 ng/mL, two-way ANOVA with Time-Combination interaction on log-transformed concentrations, Tukey HSD p=0.0001394)
d. Paxalisib mean and standard deviation brain concentration-time profiles alone (pax) and in combination (pax+mir). There are no significant differences between the groups for any time point at the a=0.05 level.

**Supplementary Fig 10: Pharmacokinetic (PK) analyses of Paxalisib and Mirdametinib show a lack of appreciable plasma or brain drug interaction and show high brain exposure**

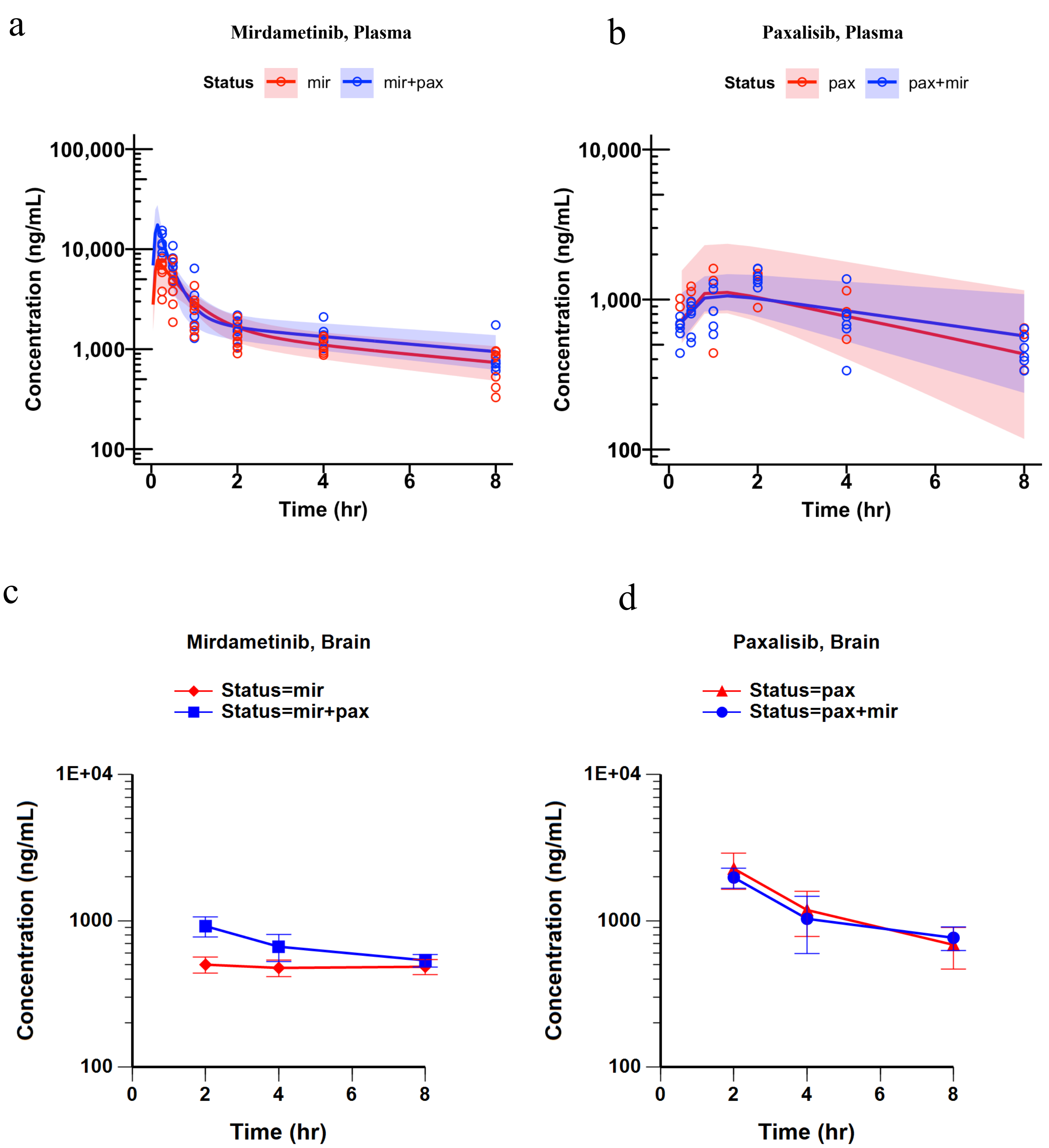

**Supplementary Figure 7: Potency and efficacy of compounds in validation screen separated by MOA.** Two-dimensional heatmaps of compounds in different MOA classes. The size of the circles is proportional to efficacy (AUC) and potency measured by compound $EC_{50}$ is depicted as a color gradient from $10^{-11}$ Molar (dark orange) to $10^{-4}$ Molar (turquoise). Rows are target class and specific compound and columns are cell models with text color coded: Tumor models are color-coded by histone H3 mutation status: H3-wt (red), H3.3G34R (blue), H3.1K27M (turquoise), and H3.3K27M (green). Control cell lines, iAstro and HABS, are black and gray, respectively. Small black dots indicate compounds not tested in that cell model. (a) The most active compounds from the screen of 93 compounds in 14 HGG cell models and two controls selected based on those inducing the top 5% AUC for at least one HGG model. All compounds evaluated in DR format are displayed by MOA in B-E and summarized in Supplementary Table 5, including (b) epigenetic modulators (c) kinases and related targets, (d) DNA damage, (e) nuclear receptors, ion channels, and G-protein coupled receptors (GPCR) and (f) compounds not covered in other mechanistic classes including ubiquitin/proteasome, heat-shock proteins, microtubules, metabolic pathways, anti-infectives, and compounds with complex MOAs.

**Supplementary Fig. 8: Pharmacodynamic analyses of mirdametinib and paxalisib show effective MEK and PI3K pathway inhibition in the brain**

Western blots with the indicated antibodies to analyze lysates from: (a) SJ-DIPGX37 intracranial tumors in mice dosed with vehicle (veh, lanes 1-3), 25 mg/kg mirdametinib (mir, lanes 4-6), or 18 mg/kg paxalisib (pax, lanes 7-9) daily for 5 days, and tissue collected 2 hours after last dose. Antibodies are shown at the right. (b) Brain from CD1-nude mice treated with vehicle (lanes 1-3), 8 mg/kg pax (lanes 4-6), 12 mg/kg pax (lanes 7-9), 8 mg/kg pax + 14 mg/kg mir (lanes 10-12) and 12 mg/kg pax + 17 mg/kg mir (lanes 13-15). (c) Brain from CD1-nude mice treated with vehicle (lanes 1-3), 14 mg/kg mir (lanes 4-6), 17 mg/kg mir (lanes 7-9), 14 mg/kg mir + 8 mg/kg pax (lanes 10-12) and 17 mg/kg mir + 12 mg/kg pax (lanes 13-15).

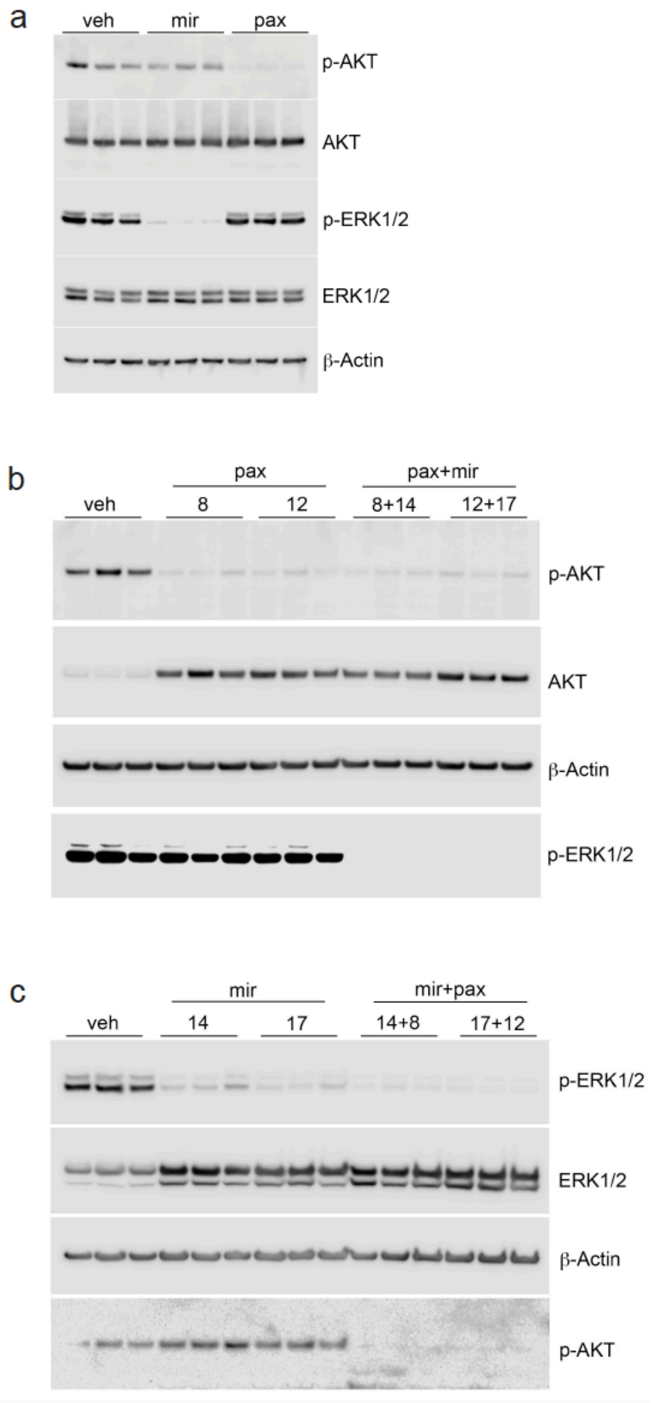

**Supplementary Fig 9: Pharmacokinetic (PK) analyses of Paxalisib and Mirdametinib show a lack of appreciable plasma or brain drug interaction and show high brain exposure**

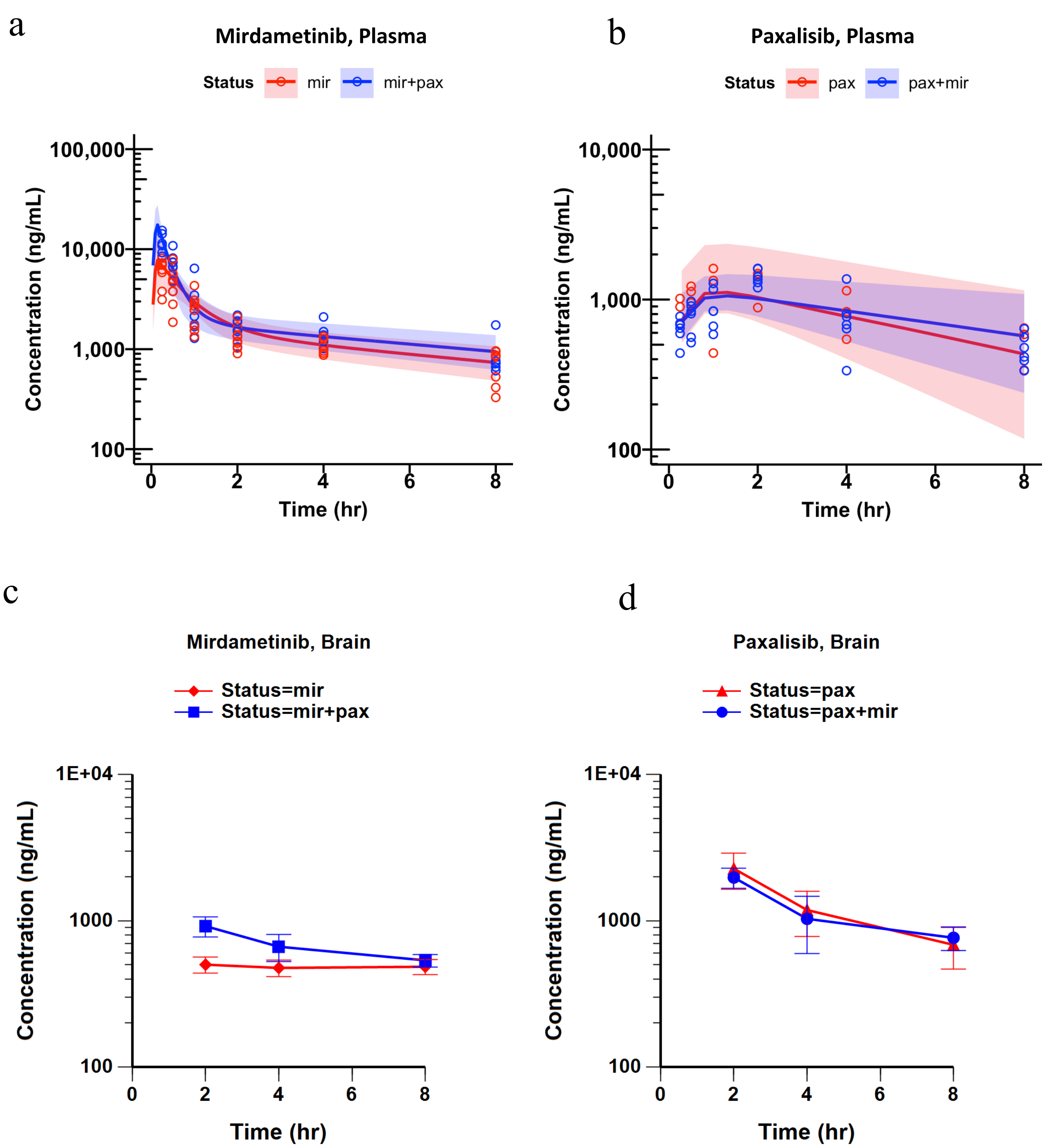

**Supplementary Fig 9: Pharmacokinetic (PK) and pharmacodynamic analyses of Paxalisib and Mirdametinib show lack of appreciable plasma or brain PK drug interaction, high brain exposure, and effective pathway inhibition**

a. Mirdametinib population mean and 90% prediction interval plasma concentration-time profiles alone (mir) and in combination with Paxalisib (mir+pax). A minor increase in Mirdametinib AUC (1.30-fold) was observed in combination.
b. Paxalisib population mean and 90% prediction interval plasma concentration-time profiles alone (pax) and in combination with Mirdametinib (pax+mir). A minor increase in Paxalisib AUC (1.63-fold) was observed in combination.
c. Mirdametinib mean and standard deviation brain concentration-time profiles alone (mir) and in combination with Paxalisib (mir+pax). Only the 2-hour time point is significantly higher for the combination (907 vs 498 ng/mL, two-way ANOVA with Time-Combination interaction on log-transformed concentrations, Tukey HSD p=0.0001394)
d. Paxalisib mean and standard deviation brain concentration-time profiles alone (pax) and in combination (pax+mir). There are no significant differences between the groups for any time point at the $\alpha$=0.05 level.

**Supplementary Tables**

**Supplementary Table 1**: **Cohort of patient tumors and derivative PDOX and cell line models.** Summary of clinical data, histopathology, DNA methylation classification and signature genes mutated for patient tumors, PDOXs, and cell lines, as well as time to PDOX engraftment.

**Supplementary Table 2**: **Sequence alterations in patient, PDOX and cell line cohorts**

(a) List of signature genes and associated pathways recurrently mutated in pHGG

(b) Non-silent mutations in HGG signature genes identified in 57 samples across 21 lines (relates to Fig. 4)

(c) Non-silent somatic mutations in non-hypermutator patient tumors, PDOXs and cell lines with matched germline available

(d) Potentially pathogenic non-silent SNVs and INDELs in patient tumors, PDOXs and cell lines without paired germline available

**Supplementary Table 3**: **GSEA of genes differentially expressed in PDOX compared with patient tumors**

(a) Genes down-regulated in PDOX samples compared to matched patient tumors (logFC < -1 & adj. P < 0.05)

(b) Significantly enriched (adj. P < 0.05) MSigDB Hallmark gene sets for the down-regulated genes in PDOX samples

(c) Genes up-regulated in PDOX samples compared to matched patient tumors (logFC > 1 & adj. P < 0.05)

(d) Significantly enriched (adj. P < 0.05) MSigDB Hallmark gene sets for the up-regulated genes in PDOX samples

**Supplementary Table 4**: **Molecular and growth characteristics of cell lines used in HTS.**

| Cell Line Name | Histone mutation status | Doubling-time* 384W (coated) (days) | Seeding number (thousand) | PDOX Methylation Classification | Signature Mutations | Reference |
|---|---|---|---|---|---|---|
| SJ-HGGX2c | Wild-type | 8.2 | 8 | Glioblastoma, IDH wildtype, subclass midline | TP53, PDGFRA, PIK3R1, MET | |
| SJ-HGGX39c | Wild-type | 3.7 | 4 | Glioblastoma, IDH wildtype, subclass RTK III | TP53, BCOR, CDKN2A, PDGFRA | |
| SJ-HGGX6c | H3.3G34R | 5.6 | 8 | Glioblastoma, IDH wildtype, H3.3 G34 mutant | H3F3A (H3.3 G34R), PDGFRA, TP53, HIST2H3D, ATRX | |
| SJ-HGGX42c | H3.3G34R | 6.1 | 8 | Glioblastoma, IDH wildtype, H3.3 G34 mutant | H3F3A (H3.3 G34R), TP53, ATRX, BCOR, PDGFRA, BRAF, CCND2 | |
| SJ-DIPGX7c | H3.3K27M | 6.1 | 8 | Diffuse midline glioma, H3 K27M mutant | H3F3A (H3.3 K27M), BCOR, TP53, PIK3CA | |
| SJ-DIPGX9c | H3.1K27M | 9.1 | 8 | Diffuse midline glioma, H3 K27M mutant | HIST1H3B (H3.1 K27M), ACVR1, PIK3CA | |
| SJ-DIPGX29c | H3.3K27M | 9.0 | 8 | Diffuse midline glioma, H3 K27M mutant | H3F3A (H3.3 K27M), TP53, CCND3 | |
| SJ-DIPGX37c | H3.3K27M | 9.0 | 8 | Diffuse midline glioma, H3 K27M mutant | H3F3A (H3.3 K27M), PIK3R1, PPM1D, NTRK1, PIK3CA | |
| SU-DIPG-IV | H3.1K27M | 3.0 | 8 | Diffuse midline glioma, H3 K27M mutant | HIST1H3B (H3.1 K27M), MDM4, ACVR1 | From Dr. M.Monje [1,2] |
| SU-DIPG-VI | H3.3K27M | 5.8 | 8 | Diffuse midline glioma, H3 K27M mutant | H3F3A (H3.3 K27M), TP53 | From Dr. M.Monje [1,2] |
| SU-DIPG-XIII | H3.3K27M | 6.6 | 8 | Diffuse midline glioma, H3 K27M mutant | H3F3A (H3.3 K27M) | From Dr. M.Monje [1,2] |
| SU-DIPG-XVII | H3.3K27M | 5.8 | 4 | Diffuse midline glioma, H3 K27M mutant | H3F3A (H3.3 K27M) | From Dr. M.Monje [2,3] |
| SUDIPG-XIX | H3.3K27M | 4.6 | 8 | Diffuse midline glioma, H3 K27M mutant | H3F3A (H3.3 K27M) | From Dr. M.Monje [2] |
| SU-DIPG-XXI | H3.1K27M | 4.6 | 8 | Diffuse midline glioma, H3 K27M mutant | HIST1H3B (H3.1 K27M) | From Dr. M.Monje [2] |
| iAstro | Wild-type | 2.8 | 4 | | | |
| HA-bs | Wild-type | 11.3 | 8 | | | |
| hNSC | Wild-type | 2.6 | 4 | | | |

*Roth V. 2006 Doubling Time Computing, Available from: http://www.doubling-time.com/compute.php

**Supplementary Table 5**: **Results of HTS.**

(a) Column definitions for DR fits for 2D vs. 3D culture condition comparison
(b) DR fits for 2D vs. 3D culture condition comparison
(c) FDA single-point screen in 9 pHGG cell line models and human embryonic stem cell-derived neural stem cells (HNSC)
(d) Column definitions for the pHGG DR screens
(e) DR fits for 246 compounds profiled in 4 exemplar pHGG cell line models
(f) DR fits for 93 compounds profiled in 14 pHGG cell line models and 2 control cell lines (iAstro and HABS)
(g) Column definitions for Combinations Studies – Raw Data
(h) Combinations Studies – Raw Data
(i) Column definitions for Combinations Studies – Fits
(j) Combination Studies - Fits

**Supplementary Table 6:** Resource Table includes all reagents used for this study

**Key Resources Table**

| | |
|---|---|
| HyClone™ Phosphate Buffered Saline (PBS) | Fisher Scientific, SH3025601 |
| Paraformaldehyde, EM Grade | Electron Microscopy Sciences, 19200 |
| Hematoxylin | Fisher Scientific, 7221 |
| Eosin | Fisher Scientific, 7111 |
| Anti-mouse biotinylated secondary antibodies | Vector Laboratories, BA-2000 |
| Anti-rabbit biotinylated secondary antibodies | Vector Laboratories, BA-1000 |
| Horseradish-peroxidase-conjugated streptavidin VECTASTAIN Elite ABC Kit | Vector Laboratories, PK-6100 |
| DAB substrates | Vector Laboratories, SK-4100 |
| Hematoxylin (for counter staining). | Vector Laboratories, H-3401 |
| RIPA Buffer (10×) | Cell Signaling, 9806 |
| Halt™ Protease Inhibitor Cocktail (100×) | ThermoFisher, 78429 |
| Halt™ Phosphatase Inhibitor Cocktail | ThermoFisher, 78420 |
| cOmplete™ Protease Inhibitor Cocktail | Sigma-Aldrich, 11697498001 |
| PhosSTOP™ | Sigma-Aldrich, 4906845001 |
| Micro BCA™ Protein Assay Kit | ThermoFisher, 23235 |
| NuPAGE MES SDS Running Buffer (20×) | ThermoFisher, NP0002 |
| NuPAGE™ 12%, Bis-Tris, 1.0 mm, Mini Protein Gel | ThermoFisher, NP0341PK2 |
| NuPAGE™ Transfer Buffer (20×) | ThermoFisher, NP00061 |
| NuPAGE™ Sample Reducing Agent (10×) | ThermoFisher, NP0004 |
| NuPAGE™ LDS Sample Buffer (4×) | ThermoFisher, NP0007 |
| NuPAGE™ 4 to 12%, Bis-Tris, 1.0 mm, Midi Protein Gel | ThermoFisher, WG1403BOX |
| Restore™ PLUS Western Blot Stripping Buffer | ThermoFisher, 46430 |
| Pierce™ 20× TBS Buffer | ThermoFisher, 28358 |
| TWEEN® 20 | Bio-Rad, 1706531 |
| Revert™ 700 Total Protein Stain for Western Blot Normalization | Li-Cor, 926-11010 |
| IRDye® 800CW Donkey anti-Rabbit IgG (H + L) | Li-Cor, 926-32213 |
| IRDye® 680RD Goat anti-Mouse IgG (H + L) | Li-Cor, 926-68070 |
| Rabbit IgG HRP Linked Whole Ab (from Donkey) | MilliporeSigma GENA934-100UL |
| SuperSignal™ West Dura Extended Duration Substrate | ThermoFisher 34075 |
| Nitrocellulose/Filter Paper Sandwiches, 0.2 µm, 8.5 x 13.5 cm | ThermoFisher LC2009 |
| Akt (pan) (C67E7) Rabbit mAb | Cell Signaling, 4691 |
| Phospho-Akt (Ser473) (D9E) XP® Rabbit mAb | Cell Signaling, 4060 |
| p44/42 MAPK (Erk1/2) Antibody | Cell Signaling, 9102 |
| Phospho-p44/42 MAPK (Erk1/2) (Thr202/Tyr204) Antibody | Cell Signaling, 9101 |

| | |
|---|---|
| Phospho-Histone H3 (Ser10) Antibody | Cell Signaling, 9701 |
| Anti-Nuclei Antibody, clone 235-1 | Millipore, MAB1281 |
| Anti-Mitochondria Antibody, surface of intact mitochondria, clone 113-1 | Millipore, MAB1273 |
| Mouse Cell Depletion Kit | Miltenyi Biotec, 130-104-694 |
| KnockOut™ DMEM/F-12 | ThermoFisher, 12660012 |
| Neurobasal® Medium, minus phenol red | ThermoFisher, 12348017 |
| B-27® Supplement, minus vitamin A | ThermoFisher, 12587010 |
| N-2 Supplement | ThermoFisher, 17502048 |
| StemPro® Neural Supplement | ThermoFisher, A1050801 |
| Recombinant Human EGF | Peprotech, AF-100-15 |
| Recombinant Human FGF-b | Peprotech, 100-18B |
| Recombinant Human PDGF-AA | Cell Guidance Systems, GFH16AF-100 |
| Recombinant Human PDGF-BB | Cell Guidance Systems, GFH18AF-100 |
| 0.2% Heparin Sodium Salt in PBS | StemCell, 07980 |
| Sodium Pyruvate (100 mM) | ThermoFisher, 11360070 |
| MEM Non-Essential Amino Acids Solution (NEAA) | ThermoFisher, 11140050 |
| GlutaMAX™ Supplement | ThermoFisher, 35050061 |
| Bovine Albumin Fraction V (7.5% solution) | ThermoFisher, 15260037 |
| Penicillin-Streptomycin (10,000 U/mL) | ThermoFisher, 15140122 |
| DMEM/F-12, HEPES | ThermoFisher, 11330032 |
| ACCUTASE™ cell detachment solution | Innovative Cell Technologies, Inc., AT 104 |
| Accumax™ cell detachment solution | Innovative Cell Technologies, Inc., AM 105 |
| HEPES (1M) | ThermoFisher, 15630080 |
| Geltrex™ LDEV-Free, hESC-Qualified | ThermoFisher, A1413302 |
| Corning™ Matrigel™ hESC-Qualified Matrix | FisherScientific,08-774-552 Corning™, 354277 |
| Primocin | Invivogen, ant-pm-1 |
| ENStem-A™ Neural Freezing Medium (1X) | EMDMillipore, SCM011 |
| STEMdiff™ Neural Progenitor Freezing Medium | StemCell, 05838 |
| Cell Freezing Medium-DMSO 1× | MilliporeSigma, C6295 |
| NeuroCult™ Proliferation Kit (Mouse & Rat) | StemCell, 05702 |
| Corning® 384-well Flat Clear Bottom White Polystyrene TC-treated Microplates | Corning, 3765 |
| BrandTech, 384-well pureGradeTM S | BrandTech, 781686 |
| Corning® 96-well polypropylene deep well V-bottom | Corning, 3960 |
| Corning® Ultra-Low Attachment 25cm² Rectangular Canted Neck Cell Culture Flask with Vent Cap | Corning, 3815 |
| Corning® Ultra-Low Attachment 75cm² U-Flask Canted Neck Cell Culture Flask with Vent Cap | Corning, 3814 |

| | |
|---|---|
| Corning® 75cm² U-Shaped Canted Neck Cell Culture Flask with Vent Cap | Corning, 430641U |
| Corning® 175cm² U-Shaped Angled Neck Cell Culture Flask with Vent Cap | Corning, 431080 |
| Falcon™ Cell Strainers | FisherScientific, 08-771-1 |
| MACS SmartStrainers (30 µm) | Miltenyi Biotec, 130-098-458 |
| MidiMACS™ Separator and Starting Kits | Miltenyi Biotec, 130-090-329 |
| LS Columns | Miltenyi Biotec, 130-042-401 |
| Papain | Worthington, LS003126 |
| N-Acetyl-L-cysteine (NAC) | Sigma-Aldrich, A9165 |
| DNase I, from bovine pancreas | Sigma-Aldrich, 11284932001 |
| D-Luciferin, Firefly | PerkinElmer, 122799 |
| CellTiter-Glo® Luminescent Cell Viability Assay | Promega, G7573 |
| Methyl cellulose | Sigma-Aldrich, M7027 |
| TWEEN® 80 | Sigma-Aldrich, P4780 |
| Distilled Water for formulation | ThermoFisher, 15230147 |
| Human iPSC-derived astrocytes, Tempo-iAstro™ | Tempo Bioscience® |
| Human Astrocytes-brain stem, HA-bs | ScienCell Research Laboratories, 1840 |
| GIBCO® Human Neural Stem Cells (H9 hESC-Derived) | Invitrogen, N7800-100 |
| GDC-0084 | Chemgood (company ID#C-11-41), CAS#1382979-94-3 |
| PD-0325901 | Chemietek (Lot#03), CAS# 391210-10-9 |

**Supplementary Table 7**: Pharmacokinetic analysis of paxalisib and mirdametinib

(a) PK study design
(b) Non-linear mixed effect (NLME) plasma PK model parameter estimates for mirdametinib, 14 mg/kg PO, QD
(c) NLME plasma PK model parameter estimates for paxalisib, 8 and 10 mg/kg PO QD

#### Supplementary Materials and Methods

##### Drug-drug interaction pharmacokinetic study of mirdametinib and paxalisib in non-tumor bearing female CD-1 nude mice

##### In Vivo Pharmacokinetic (PK) Study Design

The plasma pharmacokinetics (PK) of the MEK inhibitor mirdametinib (PD-0325901) and PI3K inhibitor paxalisib (GDC-0084) were studied to determine whether a PK drug-drug interaction (DDI) exists between these agents when co-administered orally in mice. A moderate interaction, defined as a ≥2-fold difference in plasma area under the concentration-time curve (AUC) or apparent oral clearance (CL/F), was considered as impactful and practically significant.

In the main study (Study 1), 3 groups of 9 mice each were studied with a mixed, staggered survival and terminal sampling design. Mirdametinib (Chemietech, CT-PD03, Lot# 3) and paxalisib (Chemgood, C-1141) were each suspended in 1% methylcellulose (type 400 cPs) / 1% Tween 80, with the combination co-formulated and administered as a single 10 mL/kg gavage. On Day 1, mice received single 5 mL/kg oral doses of each drug alone, with 2 retro-orbital blood samples obtained under isoflurane anesthesia, up to 8 hours post-dose. For the next 4 days, mice received daily (i.e. every 24 ± 2 hours) combination therapy or mirdametinib monotherapy. On Day 5, another PK study was performed following combination or mirdametinib monotherapy, with one survival and terminal blood sample acquired per mouse.

Paxalisib PK was also assessed in a similar combination PK DDI study with another targeted anti-cancer agent DrugX (Study 2). On Day 1, paxalisib single agent PK was studied. Then for the next 4 days, mice received daily combination therapy with DrugX or paxalisib monotherapy, with combination or single agent PK evaluated on Day 5, respectively.

For survival samples, blood from the retro-orbital plexus, 50 -100 µL, was collected into Sarstedt Minivette POCT KEDTA capillary devices. Plasma was immediately isolated and placed on dry ice until transfer to -80 °C for storage. At the terminal time points, blood was collected via cardiac puncture into a Sarstedt Microvette 500 KEDTA microtube, mice were perfused with PBS, and the brains extracted. All plasma and brain samples were stored on dry ice and transferred to -80 °C at the end of study. An outline of the studies and groups are presented in Supplementary Table 7a.

**Bioanalysis**

Plasma and brain homogenate (Dilution Factor = 6, with ultrapure water) were subjected to deproteinization and analyzed for mirdametinib and paxalisib concentrations using a qualified LC-MS/MS assay with loperamide as the internal standard. Stock and spiking solutions were prepared in methanol and used to spike matrix calibrators and quality controls. Matrix samples, 25 µL each, plus 25 µL of IS (5 ng/mL in methanol) were protein precipitated with 100 µL of acetonitrile. A 5 µL aliquot of the supernatant was injected onto a Shimadzu LC-20ADXR high performance liquid chromatography system via a Shimadzu SIL-20AC XR autosampler. The LC separation was performed using a Phenomenex Kinetex C18 (2.6 µm, 50 mm x 2.1 mm) column at 40˚C with gradient elution at a flow rate of 0.25 mL/min. The binary mobile phase consisted of 0.2% formic acid in methanol: water (10:90, v/v) in reservoir A and 0.2% formic acid in methanol in reservoir B. The initial mobile phase composition was maintained at 10% B for 0.5 minutes and was followed by a linear increase to 100% B in 2.5 minutes. The column was then rinsed for 1 minute at 100% B and then equilibrated at the initial conditions for 2 minutes for a total run time of 6 minutes. Under these conditions, mirdametinib eluded at 3.31 minutes, paxalisib at 3.00 minutes, and IS at 3.16 minutes. Analytes and IS were detected with tandem mass spectrometry using a SCIEX API 4000 in the positive ESI mode with the following mass were transitions monitored: mirdametinib 482.90 -> 249.00, paxalisib 383.30 -> 353.20, and loperamide 477.30 -> 266.10.

The method qualification and bioanalytical runs all passed acceptance criteria for non-GLP assay performance. A linear model ($1/X^2$ weighting) fit the calibrators across the 1 to 100 ng/mL range, with a correlation coefficient (R) of ≥0.9959. Sample dilution integrity was confirmed. The lower limit of quantitation (LLOQ), defined as a peak area signal-to-noise ratio of 5 or greater verses a matrix blank with IS, was 1 ng/mL for plasma and 6 ng/mL for brain homogenate secondary to the dilution. The intra-run precision and accuracy was ≤ 7.36% CV and 88.2% to 113%, respectively.

**Pharmacokinetic (PK) Analysis**

Summary statistics for mirdametinib and paxalisib concentration-time (Ct) data in plasma and brain were generated by study, occasion (Day 1 vs. Day 5), combination status, and nominal time point and presented in tabular form, along with Mean (SD) Ct profile figures, using Phoenix WinNonlin 8.1 (Certara USA, Inc., Princeton, NJ).

Plasma Ct data for mirdametinib and paxalisib were grouped by study, analyte, individual mouse, occasion, and combination status and analyzed using nonlinear mixed effect (NLME) modeling implemented in Monolix 2019R2 (Lixoft SAS, Antony, France). Parameters and the Fisher Information Matrix (FIM) were estimated using the stochastic approximation expectation maximization (SAEM) algorithm, and the final log-likelihood estimated with importance sampling, all using the default Monolix initial settings. A variety of models were fit to the Ct data, parameterized using apparent clearances or rate constants, volumes of distribution, and absorption rates as needed. These models were assessed for goodness of fit using the -2 log likelihood (-2LL) value, Akaike and Bayesian Information Criterion (AIC, BIC), visual predictive checks, plots of model individual and population predicted vs. observed data, residual plots, and the standard errors of parameter estimates. A log-normal inter-individual (omega) and inter-occasion (gamma) distribution was assumed on selected supported parameters, with only diagonal elements of parameter covariance matrices estimated. Additive and/or proportional residual error models were tested and implemented as supported. Beal's M3 method was used to handle any data that were below the LLOQ[4].

The compound combinations were tested as categorical covariates upon the mirdametinib or paxalisib supported PK parameters, primarily the apparent oral clearance (CL/F). The covariate effect was considered statistically significant if its addition reduced the -2LL by at least 3.84 units ($P < 0.05$, based on the $\chi^2$ test for the difference in the -2LL between two hierarchical models that differ by 1 degree of freedom). Additionally, Wald test P values were outputted for the interaction covariate effect levels by the software, and reported in the Results tables. Secondary PK parameters such as the maximum concentration (Cmax), time of Cmax (Tmax), area under the Ct curve (AUC), and apparent terminal half-life (T1/2) were derived from the model parameter estimates using standard formulae for the relevant compartmental model[5].

**Mirdametinib Pharmacokinetics in Mice**

The plasma PK of mirdametinib was well-described using a linear, two-compartment model with zero-order absorption. Absorption was rapid, with the Tmax generally occurring at the 0.25 hr time point. As there was no observable data in the absorption phase, the zero-order absorption rate (Tk0) was fixed to 0.125 hr. The plasma Ct profile showed a distribution phase lasting approximately 1 hr, followed by an apparent terminal phase with a half-life of ~ 4 hrs. No accumulation occurred with daily dosing for 5 days in mice. Apparent oral clearance (CL/F) was low at 12.9 mL/min/kg or ~14% of hepatic blood flow. The apparent volume of distribution was large and greater than total body water. The bioavailability of mirdametinib was not evaluated in this study, but has previously been reported to be ~30% in rats (data not shown).

A linear two-compartment model with inter-individual and inter-occasion variability on both apparent oral clearance (CL/F) and apparent oral volume of distribution (Vc/F), and proportional residual error best described the overall plasma Ct data. The precision of the variability estimates was poor likely due to model overparameterization and the small number of mice and samples available; however, the goodness of fit plots and post hoc individual visual predictive checks indicated adequate performance. Combination status was tested as a covariate on CL/F and Vc/F inter-occasion variability, and its addition on both parameters improved the model fit (-2LL P=0.0003714858), indicating a statistically significant effect of paxalisib co-administration on mirdametinib plasma PK.

There was a 23.1% reduction in mirdametinib CL/F with combination therapy (P=0.0124957), and a 61.9% decrease in Vc/F (P=0.000159031). This resulted in a 2.63-fold higher Cmax, but only a 1.30-fold increase in AUC. Therefore, no practical effect of paxalisib upon mirdametinib PK by the predefined ≥2-fold difference of AUC or CL/F criteria was observed. This lack of a practical effect is supported by comparison of Ct plots. The parameter estimates from the full mirdametinib model are presented in Supplementary Table 7b with the model predicted median, 90% prediction interval, and observed plasma mirdametinib concentrations presented in Supplementary Fig. 10a.

**Paxalisib Pharmacokinetics in Mice**

The plasma PK of paxalisib was well-described using a linear, one-compartment model with first-order absorption. Absorption rate was moderate and showed low-to-moderate variability, with the Tmax occurring at 1-2 hrs post-dose. The absorption rate appeared slower on Day 5 in combination vs. Day 1. The plasma Ct profile appeared monophasic, showing a terminal phase half-life ranging from 4.47 hrs alone to 7.28 hrs with mirdametinib. Significant accumulation occurred with daily dosing for 5 days in mice. Apparent oral clearance (CL/F) was low-to-moderate at 19.3 mL/min/kg or ~21.5% of hepatic blood flow. The apparent volume of distribution was large and greater than total body water. The bioavailability of paxalisib was not evaluated in this study, but has been reported to be high in various preclinical species.

A linear one-compartment model with inter-individual variability on first-order absorption rate (ka), apparent oral clearance (CL/F), apparent oral volume of distribution (Vc/F), and inter-occasion variability on apparent oral clearance (CL/F), and proportional residual error best described the overall plasma Ct data. The precision of some variability estimates was poor likely due to model overparameterization; however, the goodness of fit plots and post hoc individual visual predictive checks indicated adequate performance.

Combination status was tested as a covariate on CL/F inter-occasion variability, and its addition significantly improved the model fit (-2LL P=0.0005042182), suggesting a statistically significant effect of either mirdametinib and/or DrugX co-administration or study day on paxalisib plasma PK.

There was a 38.6% reduction in paxalisib CL/F in combination with mirdametinib on Day 5 (P=0.000218137), resulting in a statistically significant 1.63-fold increase in paxalisib AUC. This failed to meet the ≥2-fold difference of AUC or CL/F criteria, and therefore we conclude that there is no practical effect of mirdametinib upon paxalisib PK. This is also supported visually by comparison of Ct plots. The parameter estimates from the full paxalisib model are presented in Supplementary Table 7c with the model predicted median, 90% prediction interval, and observed plasma paxalisib concentrations presented in Supplementary Fig. 10b.

**Conclusions**

The plasma PK of mirdametinib 14 mg/kg PO in mice is similar to that previously reported with respect to the Ct curve shape. However, the AUC increased less than proportionally compared with clinically relevant doses of 0.5 and 1.5 mg/kg PO[6], suggesting saturable absorption, lower bioavailability, or higher clearance in our mice at the 14 mg/kg dose. Mirdametinib plasma PK after multiple doses appeared similar (Day 1 vs Day 5), suggesting it has time-invariant PK in mice. While a higher mirdametinib plasma Cmax was observed in combination, the AUCs were not practically different (1.30-fold) – as this failed to meet the ≥2-fold criteria, paxalisib has no practical effect upon mirdametinib PK in mice. The brain penetration of mirdametinib appeared similar alone and in combination on Day 5 (Kp,last 0.502 and 0.524, respectively), and similar to published data in mice[7]. The plasma PK of paxalisib 8 and 10 mg/kg PO in mice, after a single dose on Day 1, is similar to that reported at 25 mg/kg, assuming dose proportional PK[8].

The brain penetration of paxalisib on Day 5 in combination (Kp,last = 1.42) was similar to that reported previously (single dose Kp,6hr = 1.39)[8]. There was no difference in paxalisib brain concentration either alone or in combination with mirdametinib or DrugX. Paxalisib's plasma AUC was 1.63-fold higher than expected in combination with mirdametinib. While this difference was statistically significant, indicating a possible weak drug interaction, it failed to meet the ≥2-fold critera for a practical interaction. Therefore mirdametinib had no practical effect on the PK properties of paxalisib.

Mirdametinib and paxalisib total plasma AUCs at these dose levels in mice exceed those observed clinically in humans. A PK-guided clinically relevant dose for mirdametinib, equivalent to 4 mg PO BID in humans[9], would be 0.5 mg/kg PO BID in mice. Likewise for paxalisib, a PK-guided dose similar to 45 mg PO QD in humans[10] would be approximately 2 mg/kg to 4.5 mg/kg PO QD in mice.
